# Active learning guides automated discovery of DNA delivery via electroporation for non-model microbes

**DOI:** 10.1101/2025.11.18.689155

**Authors:** Stephanie L. Brumwell, Allison Lord, Anna D. Corts, Sierra Harken, Alexander Crits-Christoph, Mary-Anne Nguyen, Julia Leung, Kerrin Mendler, Ariela Esmurria, Charlie Gilbert, Lilan Ling, Henry H. Lee, Nili Ostrov

## Abstract

Delivery of recombinant DNA is foundational for understanding and engineering a target organism. Electroporation can be applied to any cell type, yet identification of a working protocol for new organisms remains bespoke and laborious because no systematic framework exists, and appropriate instrumentation is lacking. Here, we describe an automated high-throughput platform which uses active learning to discover electroporation protocols for non-model microbes. We first devised a 24-condition electroporation screen, based on systematic evaluation and selection of key parameters, that can be applied to any microbe. Facilitated by a custom-built fully-programmable electroporator, we successfully identified electroporation protocols for eight non-model bacteria using this screen alone. We then combined this electroporation screen with our pooled POSSUM plasmid library to simultaneously evaluate 408 experimental conditions per organism and identified both a protocol and a replicating plasmid for five non-model Proteobacteria spanning three major classes (alpha-, beta-, and gamma-). We report the first electroporation protocols for *Shewanella indica, Shewanella putrefaciens* 200, *Shewanella putrefaciens* 95, *Halomonas elongata, Piscinibacter sakaiensis,* and *Duganella zoogloeoides,* as well as multiple alternative protocols for *Shewanella amazonensis, Shewanella oneidensis, Azospirillum brasilense, Cupriavidus necator, Pseudomonas alcaliphila,* and *Escherichia coli.* Finally, we developed an active learning pipeline to guide the selection of parameters based on gathered experimental data. Using our robotic platform, we iteratively tested 538 conditions over three iterations to improve electroporation for the emerging industrial chassis *C. necator*, achieving 8.6-fold higher transformation efficiency than state of the art. This work establishes a discovery platform for DNA delivery to diverse and recalcitrant microbes that can be extended broadly to non-model organisms in our biosphere.

## Introduction

Genetic engineering methods require delivery of recombinant DNA into bacterial cells. Electroporation is a physical DNA delivery method that applies a high-voltage electric field to temporarily perforate the cell membrane and drive the entry of foreign genetic material^1–4^. Nearly all available instruments are simplified to deliver pre-programmed electric field settings optimized for specific organisms or cell types, limiting their versatility. Nevertheless, this method has been used to deliver linear DNA fragments, circular plasmids, and nucleic acid-protein complexes to enable genome engineering^5–7^ in select archaea, bacteria, yeast, fungi, algae, plants, and mammalian cells^8–12^.

Despite its wide utility, electroporation remains difficult to establish for diverse microbes. This is largely due to the lack of experimental and computational tools for systematically exploring the large parameter space of an electroporation experiment. For example, one of the few customizable instruments, the BTX Gemini X2 Electroporator, offers 4,234,500 unique combinations for electric fields composed of 1,000 voltages, 123 resistances, and 137 capacitance settings for exponential decay waveform alone^13^. Poorly defined parameters related to the preparation and recovery of electrocompetent cells, such as growth phase and cell concentration, along with the need for a functional outcome to select for successful DNA delivery, further contribute to the experimental complexity. This results in a laborious and time-consuming trial-and-error search for optimal electroporation conditions that would require hundreds of manual single-cuvette transformations to screen the vast range of parameters^14–18^. While statistical approaches like response surface methodology can identify optimal conditions^19^, this requires comprehensive sampling of the parameter space *a priori* without sequential learning or adaptive refinement. Advancements such as 96-well and microfluidic-based systems^20–24^ offer automation and multiplexing capabilities, however they can be limited by small sample volumes, incompatibility with microbial phenotypes, complicated fabrication requirements, or are ‘black box’ systems that preclude mechanistic learning or cross-platform protocol translation.

Here, we introduce automated and scalable pipelines for identifying electroporation parameters for non-model microbes (**Table 1**). We first evaluate fundamental parameters, such as cell preparation and electric field, in species from three bacterial families. We then devise an automatable 24-condition screen in a 96-well plate format, facilitated by a custom-built fully-programmable electroporator, and show its utility in identifying electroporation conditions for eight non-model bacteria. We implement a discovery pipeline by combining the 24-condition screen with delivery of a plasmid library and demonstrate simultaneous discovery of electroporation protocols and functional origins of replication (ORIs) in five non-model bacteria. Finally, we demonstrate an active learning process for electroporation parameter exploration and optimization enabled by an automated robotics pipeline and Bayesian optimizer. We demonstrate this pipeline on *Cupriavidus necator* to identify an electroporation protocol with 8.6-fold improvement over state of the art.

**Table 1.**
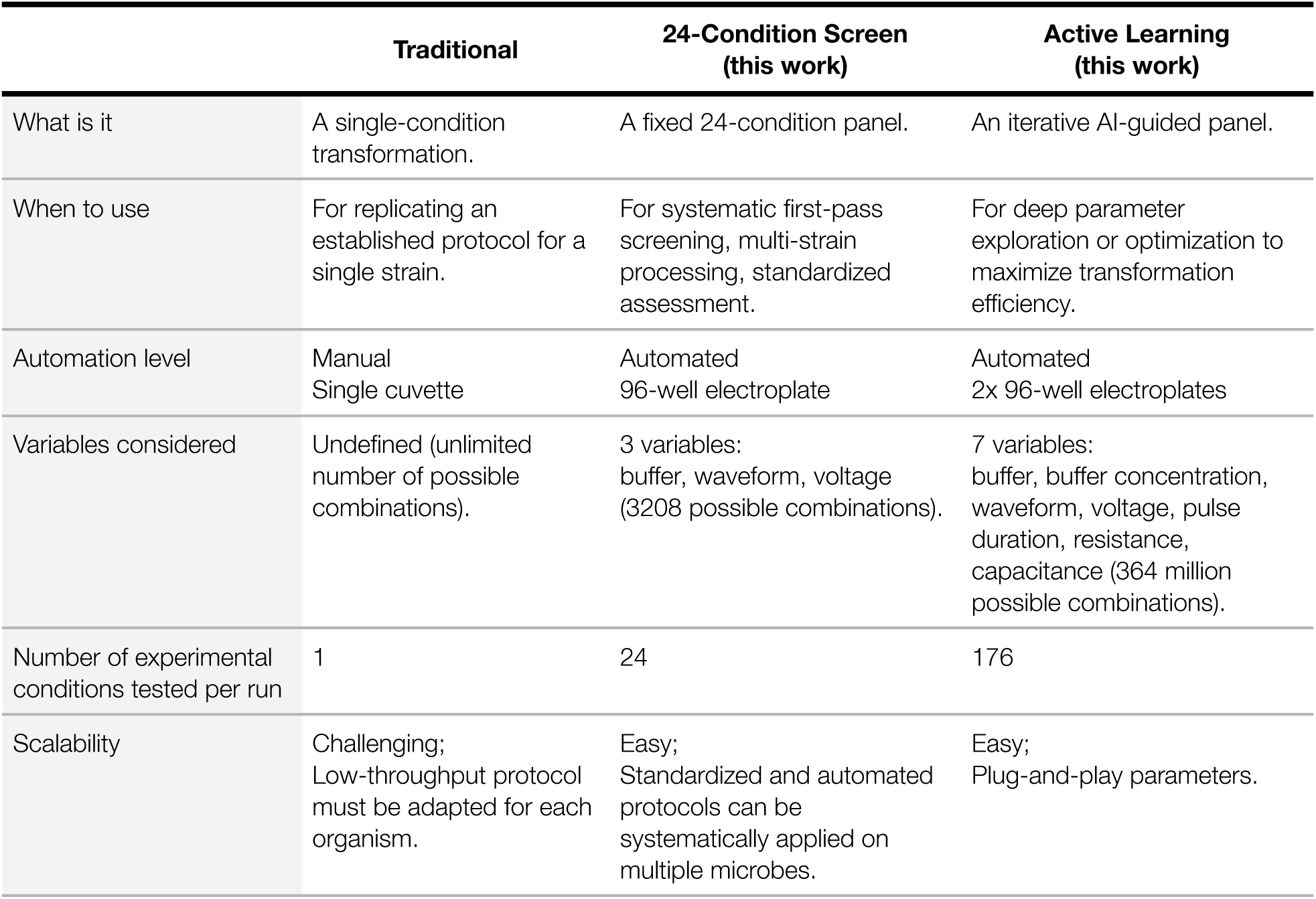
Features of the electroporation pipelines developed in this work. Comparison of utility, parameters, and execution of a traditional single-cuvette transformation to the 24-condition screen and active learning pipeline developed in this work.

|  | Traditional | 24-Condition Screen<br>(this work) | Active Learning<br>(this work) |
| --- | --- | --- | --- |
| What is it | A single-condition transformation. | A fixed 24-condition panel. | An iterative AI-guided panel. |
| When to use | For replicating an established protocol for a single strain. | For systematic first-pass screening, multi-strain processing, standardized assessment. | For deep parameter exploration or optimization to maximize transformation efficiency. |
| Automation level | Manual<br>Single cuvette | Automated<br>96-well electroplate | Automated<br>2x 96-well electroplates |
| Variables considered | Undefined (unlimited number of possible combinations). | 3 variables:<br>buffer, waveform, voltage<br>(3208 possible combinations). | 7 variables:<br>buffer, buffer concentration, waveform, voltage, pulse duration, resistance, capacitance (364 million possible combinations). |
| Number of experimental conditions tested per run | 1 | 24 | 176 |
| Scalability | Challenging;<br>Low-throughput protocol must be adapted for each organism. | Easy;<br>Standardized and automated protocols can be systematically applied on multiple microbes. | Easy;<br>Plug-and-play parameters. |

## Results and Discussion

### Evaluating key electroporation parameters

We set out to define an electroporation screen that will inform the initial parameters for various microbes, and serve as a basis for tuning an optimized high-efficiency protocol for a desired organism. To achieve this, we evaluated transformation efficiency (TE) across several commonly considered electroporation parameters, including wash buffer^25–27^ and electric field settings (waveform, voltage, resistance, and capacitance)^15–18^ (**Figure 1**). We delivered a single plasmid (**Supplementary Figure 1**, **Supplementary Table 1**) to three easy-to-culture Gram-negative bacteria with known high-efficiency electroporation protocols (10^5^–10^9^ colony forming units (CFU)/µg DNA): *Escherichia coli*^28^*, Pseudomonas alcaliphila*^29^ (relevant for wastewater bioremediation)^30^, and *Shewanella amazonensis*^31^ (relevant for microbial electrochemical systems)^3,4^ (**Supplementary Table 2-3**). All experiments were performed using 96-well electroplates (**Supplementary Figure 2**).

**Figure 1.**
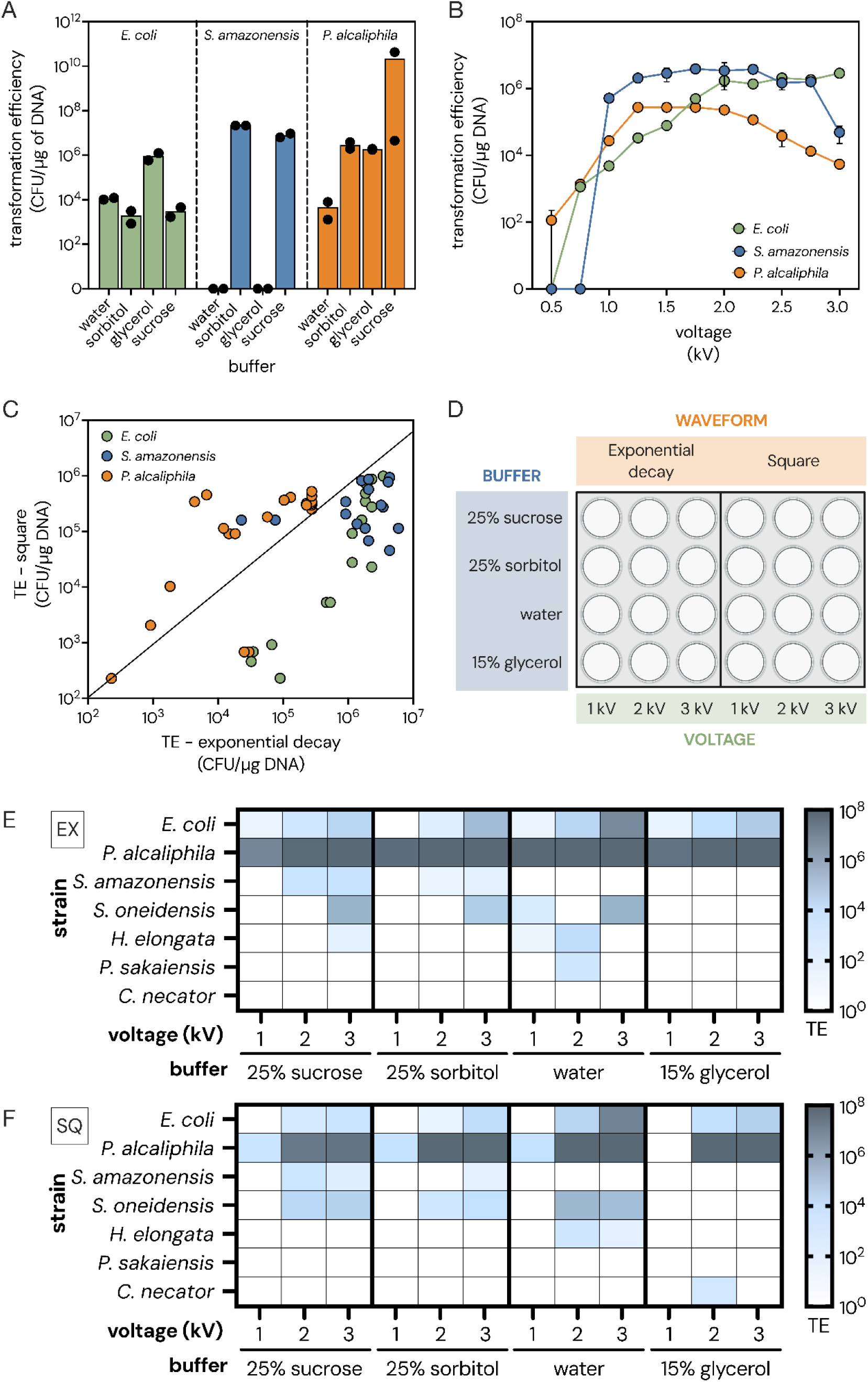
Development and application of a 24-condition electroporation screen. (**A-C**) Effect of buffer, voltage, and waveform on transformation efficiency (TE) in three Gram-negative bacteria. Data are the average of two biological replicates. (**A**) TE using four buffers: water, 25% sorbitol, 10% or 15% glycerol, and 25% sucrose. Data shown was electroporated at 3 kV using exponential decay (see additional voltages in **Supplementary** Figure 3). (**B**) TE using a range of electroporation voltages: 0.5-3 kV. Cells were washed with 10% glycerol (*E. coli*), 25% sorbitol (*S. amazonensis*), and 15% glycerol (*P. alcaliphila*), and electroporated using exponential decay waveform. Error bars represent standard error. (**C**) Comparison of TE using square or exponential decay waveforms. Data are results for all voltages and buffers tested in panel B. (**D**) Final parameter selection for 24-condition electroporation screen including four buffers, three voltages and two waveforms. (**E-F**) TE following the 24-condition electroporation screen performed on seven bacteria with a single plasmid using (**E**) exponential decay (EX) and (**F**) square (SQ) waveforms. Data are the average of two biological replicates, except *P. sakaiensis* and *C. necator* which are a single replicate.

We found that the wash buffer used to prepare cells for electroporation had a significant effect on TE, consistent with previous studies^15,17^. For each strain, one buffer showed at least 100-fold higher TE over other tested buffers (**Figure 1A**, **Supplementary Figure 3**). Using the best performing buffer for each strain, we then evaluated the effect of electric field parameters such as voltage, resistance, capacitance, and waveform type.

We found that voltage had a significant effect on TE (**Figure 1B**). We observed that 1 kV (5 kV/cm field strength) was sufficient to achieve over 10³ CFU/µg DNA across all three species. Voltages greater than 1 kV had a variable effect, resulting in up to 1000-fold improvement in some species. However, higher voltages caused a significant decrease in cell viability in all tested species (**Supplementary Figure 4**). Our results suggest resistance and capacitance have a comparatively limited effect on TE (**Supplementary Figure 5**); therefore, we set constant values for resistance (200 Ω) and capacitance (25 µF) when using exponential decay waveform.

Lastly, we investigated whether square waveforms, which are commonly limited to eukaryotic electroporation^32–34^, can be used for bacterial electroporation as they are easier to produce with modern circuits, more power efficient, and simpler to manipulate compared to exponential decay waveforms^35,36^. We found that all three bacteria could be transformed with both square and exponential decay waveforms (**Figure 1C**, **Supplementary Figure 6**). Notably, square waveform resulted in slightly increased cell survival in all organisms, particularly at higher voltages (**Supplementary Figure 6B-D**).

Based on this evaluation, we defined a minimal 24-condition electroporation screen composed of four buffers (25% sucrose, 25% sorbitol, water, and 15% glycerol), two waveforms (square and exponential decay), and three voltages (1, 2, 3 kV) (**Figure 1D**). These constrained parameters were chosen to facilitate screening of multiple diverse bacteria to identify an initial electroporation protocol, which can be optimized subsequently as desired.

### Construction of a programmable electroporator

While commercial electroporation instruments can be equipped with the use of 96-well electroporation plates, existing instruments obscure the electrical stimulus and thus scientific understanding or require human intervention to activate each electroplate column, adding unnecessary delays which reduce TE (**Supplementary Figure 7A**). To address these critical shortcomings, we custom built an electroporator that can be programmed through an application programming interface (API) and a graphical user interface (GUI). This allows for computer-controlled cycling through automated or user-defined electroporation settings for each column without the need for manual intervention for varying voltage, resistance, capacitance, pulse duration, and waveform (exponential decay and square).

### Development of an automated electroporation assay

We set out to standardize and automate our 24-condition electroporation assay so it could be applied across multiple bacteria. The first challenge we addressed was the need for highly concentrated competent cells, typically used to overcome low cell survival rates. For most microbes, obtaining highly concentrated competent cells requires processing large culture volumes that are incompatible with automation. We found that a low cell input protocol that uses unconcentrated competent cells produced a robustly detectable number of transformants (>10^2^ CFU/µg DNA) in both *S. amazonensis* and *P. alcaliphila* for the majority of voltages, albeit with lower TEs (**Supplementary Figure 8A-B**). Surprisingly, increased voltages did not significantly reduce viability of unconcentrated cells (**Supplementary Figure 8C-D**). We hypothesize that lower sample conductance reduces cell lethality compared to concentrated cultures typically used.

To compensate for reduced cell input and ensure detection of transformants when using unconcentrated cells, we extended the post-electroporation recovery period to 20 hours^15,17,18^. This change increased the detectable number of *E. coli* transformants up to 1000-fold compared with 1-hour recovery (**Supplementary Figure 7B**). When using an extended recovery period, we note that ‘transformation efficiency’ thus refers to the number of detected CFU per µg of DNA, which is *not* indicative of the number of *unique* transformation events.

Lastly, we harvested all samples at the same time regardless of culture density (OD_600_) or growth phase. While the growth phase has been shown to affect TE in some organisms^14^, a unified harvesting protocol greatly simplifies simultaneous screening of multiple non-model bacteria with different growth rates.

We used our custom electroporator to perform the 24-condition screen on nine bacteria (**Table 2**, **Figure 1E-F**, **Supplementary Figure 9-13**). For the first time, we identified electroporation protocols for *Shewanella putrefaciens* 95, *Halomonas elongata*, and *Piscinibacter sakaiensis* (formerly *Ideonella sakaiensis*). These strains represent valuable biotechnological chassis - *H. elongata* as a bioproduction host for polyhydroxyalkanoate (PHA)^37^ and ectoine^38,39^, while *P. sakaiensis* is notable for its polyethylene terephthalate (PET)-degrading capabilities^40,41^). Additionally, we identified multiple new electroporation conditions for strains with reported protocols^42–44^ including *E. coli*, *P. alcaliphila*, *S. amazonensis*, *Shewanella oneidensis,* and *Cupriavidus necator* (a bioproduction host for PHA^45–48^)) (**Figure 1E-F**, **Supplementary Figure 9A**, **Supplementary Figure 10-12**, **Supplementary Table 3**). Increasing cell concentration five-fold for *H. elongata* improved the success rate from one to six electroporation conditions (**Supplementary Figure 12B**), whereas *C. necator* showed no improvement (data not shown). We did not obtain transformants for *S. putrefaciens* 200 or *Shewanella indica,* for which no successful electroporation has been previously reported (**Supplementary Figure 9**).

**Table 2.** Highest efficiency electroporation protocols for strains screened in this study. Electroporation conditions yielding maximum transformation efficiency are reported along with the plasmid used. For strains transformed with pooled plasmid libraries, the plasmid with the highest fold-enrichment score is shown. EX, exponential decay waveform; SQ, square waveform; TE, transformation efficiency (CFU/µg DNA). Unless otherwise noted, EX resistance was 200 Ω and capacitance was 25 µF, and SQ pulse duration was 0.6 ms.

| SINGLE PLASMID DELIVERY |  |  |  |  |  |  |
| --- | --- | --- | --- | --- | --- | --- |
| Strain | Best electroporation condition (maximum TE) |  |  |  | Origin of replication | Plasmid used |
|  | TE | Buffer | Voltage (kV) | Waveform |  |  |
| *E. coli* | 10<sup>7</sup> | water | 3 | EX or SQ | pBBR1<sup>†</sup> | pAKgfp1-kan |
| *P. alcaliphila* * | 10<sup>8</sup> | 25% sorbitol | 1 | EX | pBBR1<sup>†</sup> | pAKgfp1-kan |
| *S. amazonensis* | 10<sup>4</sup> | 25% sucrose | 2 or 3 | EX | pBBR1 | pAKgfp1-kan |
| *S. oneidensis* | 10<sup>6</sup> | 25% sucrose | 3 | EX | pBBR1<sup>†</sup> | pAKgfp1-kan |
| *S. putrefaciens* 95 | 10<sup>1</sup> | 25% sorbitol | 3 | EX | pBBR1<sup>†</sup> | pAKgfp1-kan |
| *H. elongata* | 10<sup>4</sup> | water | 2 | EX | RSF1010<sup>†</sup> | pGL2_168 |
| *P. sakaiensis* | 10<sup>2</sup> | water | 2 | EX | pSa | pGL2_170 |
| *C. necator* | 10<sup>2</sup> | 15% glycerol | 2 | SQ | pSa | pGL2_270 |
| POOLED PLASMID DELIVERY |  |  |  |  |  |  |
| Strain | Best electroporation condition (maximum TE) |  |  |  | ORIs identified as functional by NGS | Plasmid with highest fold-enrichment |
|  | TE | Buffer | Voltage (kV) | Waveform |  |  |
| *S. putrefaciens* 200 | 10<sup>5</sup> | 25% sorbitol | 3 | EX | RSF1010 | pGL2_168 |
| *S. indica* | 10<sup>2</sup> | 25% sucrose | 3 | EX | RSF1010 | pGL2_168 |
| *P. alcaliphila* | 10<sup>6</sup> | water | 2 | EX | pSa<sup>†</sup>, RK2<sup>†</sup>, RSF1010<sup>†</sup> | pGL2_170 |
| *A. brasiliense* | 10<sup>6</sup> | water | 2 | EX | RK2<sup>†</sup>, pNG2 | pGL2_166 |
| *D. zoogloeoides* | 10<sup>3</sup> | 25% sucrose | 2 | EX | RSF1010, pBBR1<sup>†</sup>, pBBR1-UP<sup>†</sup>, pSa, RK2<sup>†</sup> | pGL2_168 |
| ACTIVE LEARNING |  |  |  |  |  |  |
| Strain | Best electroporation condition (maximum TE) |  |  |  | Resistance (Ω)/Capacitance (μF) | Plasmid used |
|  | TE | Buffer | Voltage (kV) | Waveform |  |  |
| *C. necator* | 10<sup>7</sup> | water | 2.555 | EX | 175 / 25 | pGL2_270 |

\* Multiple high efficiency electroporation conditions identified.
† These ORIs have been previously reported to function in the indicated species.

As part of these screens, we developed a liquid readout assay compatible with liquid-handling automation equipment. Specifically, transformants were quantified by measuring OD_600_ in liquid media. Successful transformation was evaluated by the difference in OD_600_ (ΔOD_600_) of selective media between cells electroporated with or without DNA. We compared liquid and solid selection in *E. coli*, *S. amazonensis*, *S. oneidensis,* and *H. elongata* (**Supplementary Figure 10-11**). We observed sufficient functional correlation between liquid and solid selection for both *Shewanella* species (R^2^=0.96 for *S. amazonensis* and R^2^=0.57 for *S. oneidensis*) and *H. elongata* (R^2^=0.52), but not for *E. coli* (R^2^=0.01) (**Supplementary Figure 11D**, **Supplementary Figure 12D**). Lack of correlation for *E. coli* was likely due to its faster doubling time (20 minutes compared to ∼1-2 hours for the other strains), allowing *E. coli* transformants to grow well into the stationary phase over the 24-hour selection period. Measuring OD_600_ at an earlier time or at multiple time points during liquid selection could help discriminate top-performing electroporation conditions in faster-growing organisms. We note that in some cases liquid selection identified fewer successful electroporation conditions compared to solid selection. For instance, in *S. amazonensis*, agar selection identified seven positive conditions while liquid selection identified four, missing the three conditions with the lowest reported TE (**Supplementary Figure 10C**, **Supplementary Figure 11B**). Further work is needed to investigate the cause of these differences, which may include a strain’s ability to grow in liquid versus solid media, the effective stringencies of antibiotic selection in either medium, or stress responses that prolong lag phase. Overall, we conclude that liquid selection is a sufficient and automation-compatible method to detect transformants.

These results demonstrate that the 24-condition electroporation screen can recapitulate established protocols and identify new electroporation conditions in non-model microbes. However, our inability to identify protocols for all strains exemplifies the difficulty of applying a unified protocol across organisms with diverse phenotypes. More work is needed to investigate the factors underlying variability in efficiency, which could include differences in restriction modification systems, viability in suspension buffers, growth phase, or membrane permeability^49^.

### Simultaneous discovery of electroporation protocols and functional ORIs

When attempts to deliver DNA to non-model microbes fail, it is difficult to determine whether the lack of transformants is due to an unsuitable electroporation protocol or incompatible genetic parts. Thus, the ability to simultaneously discover electroporation protocols and functional plasmids would be desirable to establish DNA delivery in non-model microbes (**Figure 2A**).

**Figure 2.**
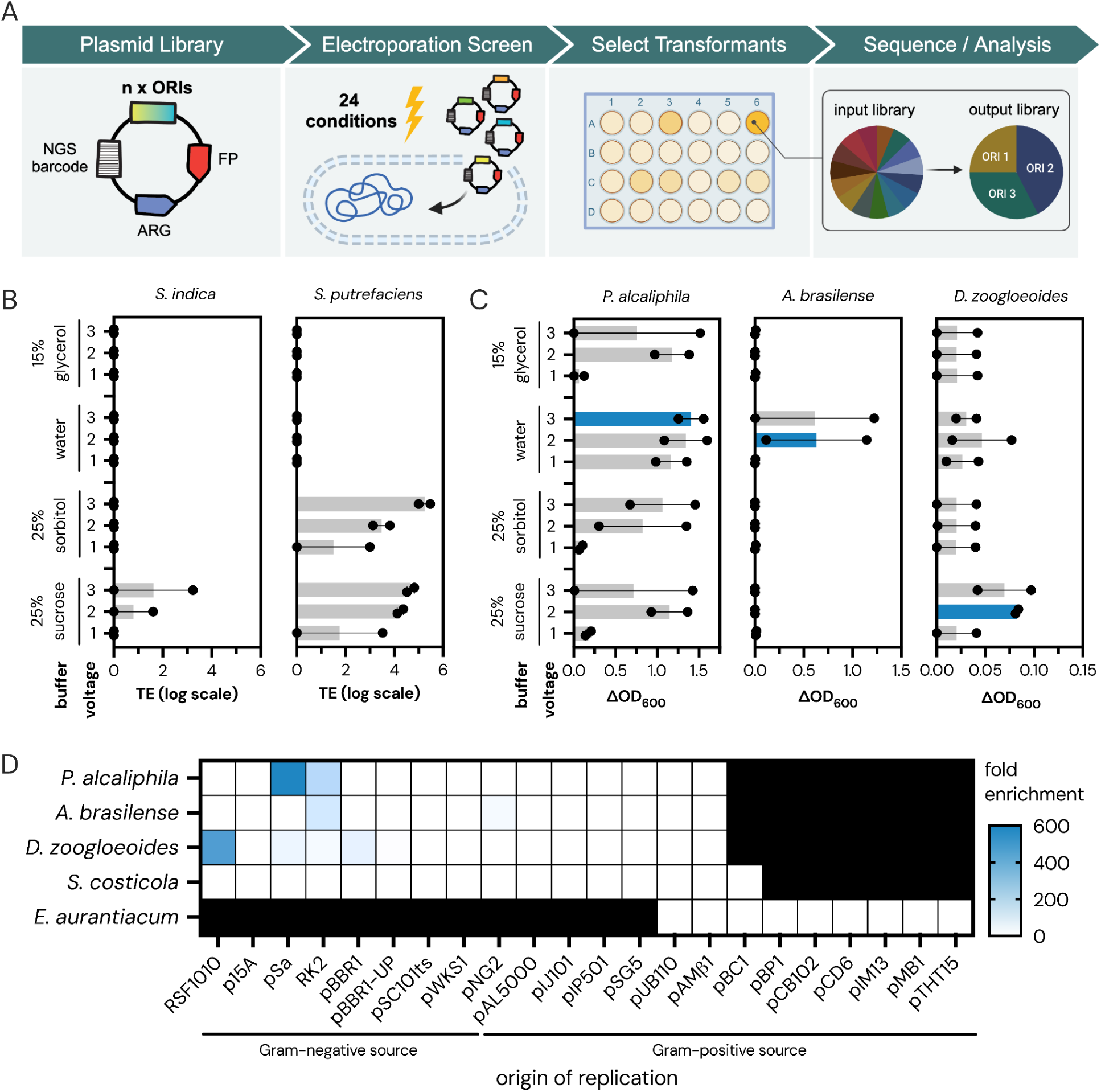
Discovery of electroporation conditions and functional origins of replication in five bacteria. (**A**) Schematic of pooled plasmid library 24-condition electroporation screen. First, plasmids harboring different ORIs and unique barcodes for downstream analysis are pooled. Then, the pooled plasmid library is delivered to target bacteria using the 24-condition electroporation screen. Next, transformants are selected in liquid media and on agar to determine viable electroporation conditions. Finally, amplicon sequencing of the plasmid barcodes is used to identify functional ORIs in each strain. FP, fluorescent protein; ARG, antibiotic resistance gene. (**B**) Transformation efficiency (TE) of a four-plasmid pooled library in *S. indica* and *S. putrefaciens* 200. Data is shown for exponential decay using solid selection only. Data are the average of two biological replicates. Error bars represent standard error. (**C**) Results of electroporation screen using a pooled plasmid library in *P. alcaliphila, A. brasilense, D. zoogloeoides, S. costicola,* and *E. aurantiacum.* Data is shown for exponential decay using liquid selection only, as measured by ΔOD_600_. See **Supplementary** Figure 14**-15** for full results. The blue bar indicates the electroporation condition with the highest ΔOD_600_ that was chosen for downstream sequencing analysis. Data are the average of two biological replicates. Error bars represent standard error. (**D**) ORIs identified in each bacteria by sequencing transformants from the best electroporation condition (highest ΔOD_600_) as shown in panel C. Color gradient represents fold enrichment of each ORI over control (non-functional dummy ORI). Black boxes indicate that the plasmid was not included in the input library. White boxes indicate that the ORI was not detected in the sequenced sample. Data are the average of two biological replicates.

We first tested the feasibility of pairing our 24-condition screen with a small four-plasmid ORI-variant library (RSF1010, RK2, pNG2, and p15A) in the two *Shewanella* strains that failed previous electroporation attempts, *S. indica* and *S. putrefaciens* 200 (**Supplementary Table 1**, **Supplementary Figure 9**). Encouragingly, we were able to identify successful transformation conditions using the 4-member pooled plasmid library (**Figure 2B**, **Table 2**). Sequencing of transformants revealed RSF1010 as the ORI driving replication in both strains. In *S. putrefaciens* 200 we obtained 10^2^–10^6^ CFU/µg DNA using sucrose or sorbitol and either exponential decay or square waveforms. *S. indica* yielded a lower efficiency (up to ∼10^2^ CFU/µg DNA) using sucrose or sorbitol, limited only to exponential decay. Neither species yielded transformants with water or glycerol, consistent with previous results obtained for other *Shewanella* species.

Using an expanded number of plasmids from our POSSUM toolkit^50^ (10- or 17-plasmid libraries) we were able to identify both electroporation conditions and functional ORIs in three Gram-negative non-model bacteria (**Figure 2C**, **Table 2**, **Supplementary Figure 14-15**). Specifically, this pipeline identified six electroporation conditions for the antifungal producer *Duganella zoogloeoides* (for which no known protocol exists), 13 electroporation conditions for the plant-associated bacterium *Azospirillum brasilense*, and many conditions for *P. alcaliphila*. Notably, the same high-efficiency conditions were identified in both liquid and agar selection (**Supplementary Figure 14-15**). No transformants were observed for extremophiles *Salinivibrio costicola* (Gram-negative) and *Exiguobacterium aurantiacum* (Gram-positive).

In each library experiment, we extracted plasmid DNA from transformants and sequenced the DNA barcodes encoded on the plasmids^43^ to identify functional ORIs in each electroporation condition (**Supplementary Table 4**). In *P. alcaliphila,* we recovered plasmid ORIs pSa and RK2 from all samples that yielded significant liquid growth (ΔOD_600_ >0.25) (**Figure 2D**, **Supplementary Figure 16**), and RSF1010 in three of the sequenced samples. These results align with our findings through conjugation experiments in *P. alcaliphila*^43^ and previous studies^29^. In *A. brasilense* we identified the previously reported RK2^51^ ORI and newly discovered pNG2. In *D. zoogloeoides,* we identified five functional ORIs including two novel (RSF1010 and RK2) and three previously reported using conjugation (pBBR1, pBBR1-UP, and RK2)^43,52^ (**Figure 2D**). However, our library screen failed to detect some functional ORIs previously indicated through single-plasmid electroporation or conjugation, such as ORI pBBR1 in *P. alcaliphila* (**Figure 1**), ORI p15A in *D. zoogloeoides*^43,52^, or functional ORIs in *S. costicola* and *E. aurantiacum* (data not shown)^53,54^. This may be due to the smaller dynamic range of a pooled library, especially if competition between bacteria harboring different plasmid ORIs result in loss of ORIs from the population^55^. We hypothesize that restriction-modification systems may be the reason for ORIs found through conjugation but not electroporation, as conjugation has been postulated to evade several restriction systems through single-stranded DNA delivery^56^. Further work to test this is underway.

Increasing the number of ORIs in our plasmid libraries will broaden the range of bacteria that can be addressed. While our libraries already include many ORIs used in laboratories to date^50^, we are continuing to expand this collection with additional natural plasmids which may have narrow or broad host range^57^. These plasmids can be recoded to evade innate bacterial defense systems. Even if a functional ORI is present, the host may digest the incoming plasmid DNA with defense systems such as restriction-modification, which can be detected with Nanopore sequencing^58,59^ and specific sequences can be protected^60–63^ or removed^64,65^ to evade degradation. However, the full array of bacterial defense systems are still being uncovered^66^. To counteract these systems, anti-defense systems^67^ may be added to our plasmid libraries.

### Development of an electroporation active learning pipeline

An automated robotics pipeline can enable new approaches in high-throughput exploration and optimization of electroporation protocols. For many organisms with no published protocols, it is difficult to know which parameters to start with and iterate upon. We demonstrate an active learning process to guide parameter selection for optimization of DNA delivery in new organisms. An iterative Bayesian optimization process synthesizes cumulative data generated in each experimental cycle to identify optimal electroporation conditions that maximize TE for a given strain (**Figure 3A**).

**Figure 3.**
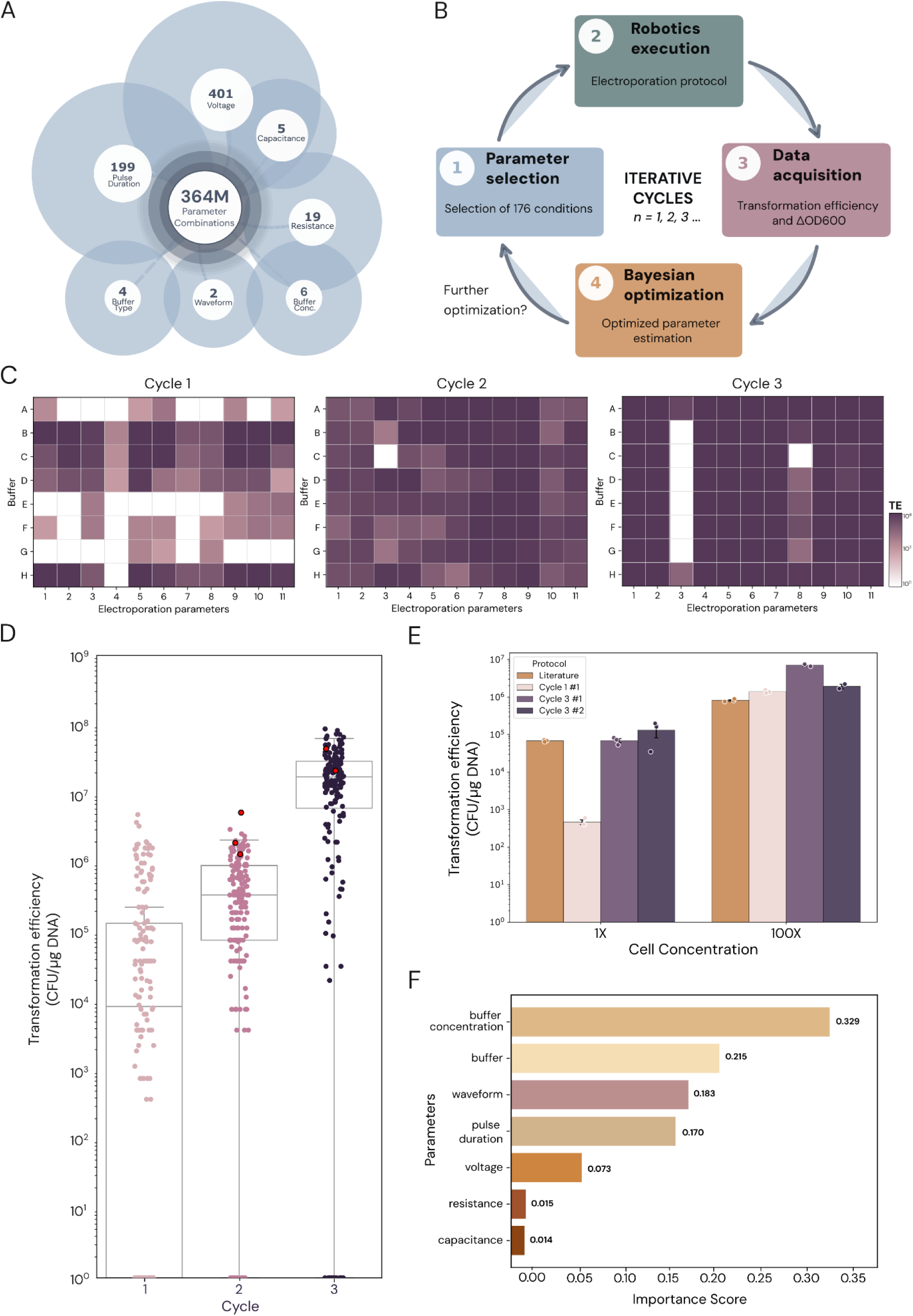
Electroporation active learning pipeline applied to *Cupriavidus necator*. (**A**) Description of the electroporation parameter space explored in this study. (**B**) A diagram of the active learning process. (**C**) Electroplate map displaying transformation efficiency (CFU/µg DNA) of one of the electroplates (88 out of 176 conditions) from each cycle of active learning. (**D**) Transformation efficiency (CFU/µg DNA) for 176 samples tested in each active learning cycle. The red points represent the best performing parameter from Cycle 2 repeated in Cycle 3. (**E**) Transformation efficiency of unconcentrated (1X) and concentrated (100X) cells using the best electroporation protocol from Cycle 1, the two best protocols from Cycle 3, and the current gold standard from literature^68^. Error bars represent standard error of the mean. (**F**) Importance scores for each electroporation parameter’s impact on TE optimization, computed using functional ANOVA.

In each experimental cycle, we used Bayesian optimization to select up to 176 combinations of electroporation parameters from a 364-million electroporation parameter space for buffer type, buffer concentration, waveform, voltage, pulse duration^16,32–34^ (square wave), resistance and capacitance (exponential decay) (**Supplementary Table 5**). We used our automated pipeline to execute the recommended experiments and feed the results back into the Bayesian optimizer to select parameters for the next iteration (**Figure 3B**).

We demonstrated this pipeline on the Gram-negative bacterium *Cupriavidus necator* (formerly *Ralstonia eutropha*), industrially relevant for CO_2_ valorization and production of bioplastics (PHAs)^45–48^. We tested if our active learning approach could identify a wild-type *C. necator* electroporation protocol equivalent or superior to state of the art published protocols, which increased efficiency from 10^5^ to 10^8^ CFU/µg DNA.

We performed three cycles of active learning and optimization (**Figure 3C-D**). Each cycle consisted of parameter suggestions followed by experimental electroporation using an automated robotic pipeline. In the first cycle, 176 space-filling and uniform selected conditions were determined by a Quasi Monte Carlo sampler algorithm for all seven parameters without using preexisting data. We then tested all conditions in parallel using our automated electroporation pipeline. While 33% (58 conditions) failed to produce transformants, we observed that 67% (118 conditions) produced transformants with a TE of 10^2^-10^6^. Based on processing of Cycle 1 results, the algorithm then suggested 176 additional conditions that were executed in the second cycle of experiments. The mean of all Cycle 2 samples was 1.95-fold higher than Cycle 1, with 12 conditions failing to produce transformants. The number of successful conditions increased to 93%, with only 12 conditions failing to produce any transformants, and the mean efficiency improved by about two-fold. In the third and final cycle, we again observed an increase in the number of successful protocols as well as the overall efficiency across all experiments. The mean of all Cycle 3 samples was 34.6-fold higher than Cycle 2, with TE ranging from 10^4^-10^7^, or 3-fold higher after normalizing for batch effects by including the previous best performing conditions from Cycle 2 in Cycle 3 (**Figure 3D**). We found good correlation between liquid and solid selection results, indicating either selective strategy could be used (**Supplementary Figure 17**).

With successive cycles, the Bayesian optimizer narrowed in on successful parameters (**Supplementary Figure 18**). To assess the contribution of each electroporation parameter to the overall efficiency, we applied functional ANOVA importance evaluation^69^. In the three cycles performed for *C. necator*, the algorithm identified buffer concentration, buffer type, and waveform as the most important parameters for optimizing efficiency, while resistance and capacitance had the lowest importance scores (**Figure 3F**). In sum, we show that a Bayesian optimization approach combined with a robotic experimentation platform successfully identified a superior electroporation protocol for an emerging industrial microbial chassis within three weeks.

To validate our results, we compared the best condition identified from Cycle 1, Cycle 3, and the optimized published protocol for *C. necator*^68^ (**Figure 3E**, **Table 2**). This experiment was performed as published: manually using a single-sample cuvette, concentrated cells, and 2-hour recovery. Using this protocol, we found the top-performing Cycle 3 protocol resulted in 8.6-fold higher TE than the literature protocol and 5-fold improvement compared to the Cycle 1 protocol (**Figure 3D-E**). Notably, even when using unconcentrated cells as we do in our automated pipeline, the top performing Cycle 3 protocol was 200-fold better than Cycle 1 hits and similar to the optimized published protocol. While this underscores the effect of concentrated cells on efficiency, it also highlights the feasibility of using the easier, faster non-concentrated preparation of cells to explore and optimize electroporation conditions. Following three cycles of active learning, the protocol in our assay outperformed all previously tested conditions as well as the published optimized protocol (**Figure 3E**).

Active learning now enables resource-efficient testing of parameters for a given non-model microbe. For example, additional buffer compositions and chemistries could be explored beyond the four buffers presented in this work. Altering growth^15^ or recovery medium^18^ can positively affect membrane composition and mitigate stress or defense systems. Harvesting cells in the exponential or stationary growth phase may impact TE^18^. It may also be relevant to alter the temperatures used for the preparation of competent cells or incubation in the cuvette^15^, recovery conditions, or input plasmid concentration^16,55^. Improvements may be gained by weakening the cell wall through chemical or enzymatic^70^ methods, particularly for Gram-positive bacteria due to their thicker peptidoglycan layer^17^. Finally, additional custom electric field variations may be tried, such as pulse polarity, number of pulses and alternative waveforms such as sinusoidal or triangle.

## Conclusion

To date, electroporation protocols require empirical optimization across a vast, multidimensional parameter space with no systematic framework or predictive models. By building an automated platform with custom instrumentation and robotic integration, it became feasible to systematically study this parameter space. To quickly establish initial electroporation conditions for a new bacteria, we devised a compact 24-condition assay and showed its utility for 12 bacterial species using both single plasmids and pooled plasmid libraries. By integrating active learning, we show that this platform can now iteratively improve initial conditions to drive towards optimal parameters by demonstrating an improvement over state of the art delivery for *C. necator*. This automated system makes it possible to iteratively test a large range of parameters relevant to diverse organisms, including yeasts, fungi, archaea, and other non-model eukaryotes.

Broadly, we demonstrate the use of bespoke laboratory instrumentation and robotic automation to address longstanding challenges for biological research to systematically collect data for predictive models. With ongoing integration of additional artificial intelligence advances, including model context protocol (MCP) for aggregating information and agents for reasoning and data analyses, this platform serves as a biological research co-pilot for scientists and paves the way for fully self-driving autonomous laboratories.

## Methods

### Microbial strains and growth conditions

Bacterial strains used in the study are summarized in **Supplementary Table 2**. Bacterial growth conditions for all strains including media, growth temperature, and antibiotic concentrations used for media supplementation are summarized in **Supplementary Table 6**. Solid media was made using 1.5% agar. Strains grown in liquid media in flasks or conical tubes were shaken at 225 RPM, while strains grown in 2-mL 96-deep well plates with pyramid bottoms (Agilent or Nest) were shaken at 800 RPM on a digital microplate shaker (Thermo Fisher Scientific).

### pAKgfp1-kan plasmid construction

Plasmids used in this study are summarized in **Supplementary Table 1**. Gibson Assembly was used to modify pAKgfp1^71^, replacing the ampicillin selective marker with the kanamycin selective marker from pGLDV_5 [28o1G]^43^. To achieve this, Q5 High-Fidelity 2X Master Mix (New England Biolabs) was used to amplify the pAKgfp1 backbone (∼4.8 kbp) and kanamycin marker (∼0.9 kbp). The PCR parameters used to amplify each fragment are specified below and the primers used are listed in **Supplementary Table 4**.

<u>PCR.</u> The backbone was amplified in a 50 µL reaction prepared according to the manufacturer’s instructions, using pAKgfp1^71^ as template DNA. PCR was performed using the following thermocycler conditions: 98°C for 30 sec, 35 cycles of 98°C denaturation for 10 sec, 58°C annealing for 15 sec, 72°C extension for 2.5 min, and a final 72°C extension for 2 min. The kanamycin marker was amplified in a 25 µL reaction prepared according to the manufacturer’s instructions, with 2 µL of 5–8 ng/µL pGLDV_5 [28o1G]^43^ template DNA. PCR was performed using the following thermocycler conditions: 98°C for 30 sec, 30 cycles of 98°C denaturation for 10 sec, 61°C annealing for 15 sec, 72°C extension for 30 sec, and a final 72°C extension for 2 min. The products of both PCR reactions were DpnI digested for 1 hour and gel-purified using the DNA Clean & Concentrator-5 kit (Zymo Research).

<u>Assembly.</u> The final plasmid was assembled from these two PCR fragments in a 20 µL Gibson Assembly reaction using NEBuilder HiFi DNA Assembly Master Mix (New England Biolabs) according to the manufacturer’s instructions, using 1:2 vector to insert ratio (50 ng vector:100 ng insert). The assembly mix was incubated on a thermocycler for 15 minutes and then stored at –20°C. The final plasmid, pAKgfp1-kan, was transformed into NEB 5-alpha electrocompetent *E. coli* (New England Biolabs), extracted by plasmid miniprep using the QIAprep Spin Miniprep Kit (Qiagen), and sequence verified. The plasmid was then transformed into *dam–/dcm–*electrocompetent *E. coli* (New England Biolabs) according to the manufacturer’s instructions, extracted and sequence verified by Eton Bioscience (Boston, MA).

### Plasmid DNA extraction

Individual pGL2 plasmids (**Supplementary Table 1**) were stored in TransforMax EC100D pir-116 cells (Biosearch Technologies); pAKgfp1-kan was stored in *dam–/dcm– E. coli* (New England Biolabs). *E. coli* strains harboring the pGL2 or pAKgfp1-kan plasmids were grown overnight shaking in LB media supplemented with the appropriate antibiotic (gentamicin, kanamycin or chloramphenicol). Plasmids were extracted from saturated cultures using the QIAprep Spin Miniprep Kit (Qiagen, CAT#27104), eluted in nuclease-free water to avoid arcing, and stored at −20°C. For large quantities, Gigaprep plasmid preparation was performed by Eton Bioscience (Boston, MA). DNA concentrations were quantified with a Qubit 1X dsDNA High Sensitivity Kit (Invitrogen) and adjusted to ∼100 ng/µL with nuclease-free water.

### POSSUM Toolkit plasmid library preparation

Plasmid libraries used in this study are summarized in **Supplementary Table 1**. <u>Cloning</u>. The 10 plasmids included in the chloramphenicol plasmid library were built using parts from the POSSUM Toolkit^50^ (**Supplementary Table 7**) and assembled using Loop Assembly as previously described^50^. The plasmids included in the 17-plasmid gentamicin library were previously constructed using this same method^50^. <u>Pooling</u>. The extracted pGL2 plasmids^50^ included in the 4-, 10- and 17-plasmid libraries were pooled in equal concentration and adjusted with nuclease-free water to achieve a final, total DNA concentration of 100 ng/µL.

### Electroporation buffer preparation

Buffers used in the parameter screen and 24-condition screen include 25% sucrose (304.3 mL/L 1.2 M sucrose (Boston Bioproducts, CAT#IBB-255)), 25% sorbitol (250 g/L D-sorbitol (Fisher Chemical, CAT#S459-500)), water (Milli-Q), and 10% or 15% glycerol (200 or 300 mL/L 50% glycerol (Teknova, CAT#G1798)). All buffers were dissolved in Milli-Q water to a final volume of 1 L and filter-sterilized with 0.2 µm PES filters (Thermo Fisher Scientific). For active learning experiments, sucrose, sorbitol, and glycerol buffers were made in a range of concentrations from 5-30% in 5% increments from a 30% stock solution of each buffer.

### Electroporation protocol - parameter screen

Unless otherwise stated, all protocol steps were performed at room temperature, which simplifies future automation and has been shown to improve transformation and editing efficiency in previous studies^44,72^.

<u>Culturing.</u> Strains were struck out from 20% glycerol stocks onto LB agar and incubated at optimal growth temperature until colonies appeared. Cultures were inoculated with two single colonies from <2-week-old streak plates into two baffled flasks containing 20–50 mL of LB broth. Cultures were incubated overnight (16–18 hours) shaking at optimal growth temperature. Overnight cultures were diluted into 50–100 mL of LB media in two or more flasks (depending on the scale of the experiment performed) to an OD_600_ of 0.08, then grown in the same conditions to an OD_600_ defined in previous studies (∼1-2 for *S. amazonensis*^31^ and *P. alcaliphila*^29^, or ∼0.4 for *E. coli*^28^).

<u>Competent cell preparation.</u> Cultures were pelleted by centrifugation in 50-mL centrifuge bottles at 5000 RCF for 20 minutes at 4°C. Supernatants were carefully discarded, and the pellet was gently resuspended by pipetting in a volume of the appropriate wash buffer equivalent to a third of the subculture volume. This process was repeated for a total of three washes. After the third wash, the supernatant was discarded and a volume of wash buffer was added to concentrate cells 100-fold the initial culture volume for *E. coli* and *P. alcaliphila,* or 60-fold for *S. amazonensis*^31^.

<u>Electroporation.</u> Unless otherwise stated, all electroporations were performed in a 0.2-cm gap electroplate or cuvette (as indicated). For each transformation, 5 μL of ∼100 ng/µL plasmid DNA was added to each well, followed by 95 μL of competent cells, and mixed by gentle pipetting three times. For negative controls, sterile nuclease-free water was used instead of DNA and electroporated using exponential decay and 1 kV. Unused electroplate wells in the same columns were filled with 100 μL of wash buffer. Exponential decay protocols were performed with a resistance of 200 Ω and capacitance of 25 µF. Square wave protocols were performed with a 0.6 ms pulse duration. Cuvette transformations were performed with a BTX Gemini X2 twin-wave electroporator. Electroplate transformations were performed using our custom electroporator.

<u>Recovery.</u> Electroplate samples were recovered by quickly transferring transformed cells (100 μL) into 900 μL of LB broth in a 96-deep well plate with a multichannel pipette. Cuvette samples were recovered by adding 900 μL of LB directly into the cuvette and transferring mixture into a 96-deep well plate for incubation. Samples were recovered by shaking at 1000 RPM at 37°C. Strain-dependent recovery time was chosen to prevent doubling: *E. coli,* 1 hour; *P. alcaliphila*, 1.5 hours; and *S. amazonensis*, 2 hours.

<u>Selection</u>. After recovery, the samples were serially-diluted 10-fold (100 µL into 900 µL of LB) up to 10^-8^ and spot-plated on selective and non-selective media. The 10^-4^–10^-8^ dilutions were plated on nonselective media to score surviving cells. The 10^0-4^ dilutions were plated on selective media supplemented with the appropriate antibiotic to score transformants. Plates were incubated at the optimal growth temperature for each strain (**Supplementary Table 6**) for 1–2 days until colonies appeared. Colonies were counted manually.

### Electroplate cleaning

Following electroporation, electroplates were washed according to the manufacturer’s instructions (BTX)^73^ and were reused. Briefly, the wells were rinsed i) three times with filter-sterilized (0.2 µm PES filter, Thermo Fisher Scientific) Milli-Q water, ii) once with 100% ethanol, and iii) once with 70% ethanol, and left open to dry overnight in a biosafety cabinet. The dried electroplates were sealed with sealing tape (Thermo Fisher Scientific) and stored in a sterile plastic bag at room temperature.

### Electroporation protocol - 24-condition screen

Unless otherwise stated, all protocol steps were performed at room temperature. General experimental steps are outlined in **Supplementary Table 8**.

<u>Culturing</u>. Initial cultures were inoculated directly from glycerol stocks into 1 mL of appropriate growth media in a 96-deep well plate, sealed with a breathable membrane (ThermoFisher), and incubated with shaking for 24 hours. Cultures were then pelleted by centrifugation at 5000 RCF for 5 minutes at room temperature (Beckman Allegra X-12 Centrifuge) and resuspended in 1 mL of media. The washed culture was diluted into four wells of a 96-deep well plate containing growth media to a final volume of 1 mL. The dilution ratio was determined by the doubling time of the strain (<2.5 hours = 1:50, >2.5 hours = 1:10). The 96-deep well plate was sealed with a breathable membrane and incubated with shaking for 4 hours. Note: i) For *P. sakaiensis*, an initial 40-mL culture was grown and 1 mL volumes were aliquoted into the 96-deep well plate to be processed as described; ii) In some cases, multiple dilution ratios were attempted for the same strain.

<u>Competent cell preparation.</u> After 4 hours, cells were pelleted by centrifugation at 5000 RCF for 5 minutes at room temperature. Using the CyBio FeliX or Opentrons Flex, ∼950 μL of supernatant was removed. Cells were washed with 700 µL of the appropriate wash buffer (differing by row). Centrifugation and wash steps were repeated for a total of four washes but in subsequent wash steps ∼700 μL of supernatant was removed. This yielded ∼800 μL of unconcentrated competent cells, sufficient for seven transformation reactions.

<u>Electroporation.</u> All electroporations were performed in a 0.2-cm gap electroplate using our custom electroporator. For each transformation, 5 μL of ∼100 ng/µL plasmid DNA was added to each well, followed by 95 μL of competent cells. For negative controls, no DNA was added and cells were electroporated using exponential decay and 1 kV. Unused electroplate wells in the same columns were filled with 100 μL of wash buffer. Exponential decay protocols were performed with a resistance of 200 Ω and capacitance of 25 µF. Square wave protocols were performed with a 0.6 ms pulse duration.

<u>Recovery.</u> Electroplate samples were recovered using the CyBio FeliX to quickly transfer transformed cells (100 μL) into 900 μL of growth media in a 96-deep well plate. The plate was sealed with a breathable membrane and samples were recovered for 20 hours by shaking at 800 RPM at the optimal growth temperature (**Supplementary Table 6**).

<u>Selection</u>. After recovery, transformants were selected on agar and/or in liquid media. The samples were serially-diluted 10-fold (100 µL into 900 µL of PBS) up to 10^-2^ using the CyBio FeliX or Opentrons Flex. For solid selection, 5 μL of the 10^0^-10^-2^ plates was spot plated on selective and nonselective media using the Nayo N96. Plates were incubated at the optimal growth temperature for each strain until colonies appeared. For liquid selection, 50 μL from the 10^0^-10^-2^ plates was inoculated into 950 µL of selective and nonselective media in a 96-deep well plate using the CyBio FeliX or Opentrons Flex and incubated at optimal growth temperature with shaking for 24 hours (or until growth was seen).

<u>Data collection</u>. Colonies on solid plates were counted manually. OD_600_ of liquid cultures were measured using 100 µL of sample on the Synergy H1 plate reader.

### Electroporation protocol - four-plasmid library

All steps were performed as described in the ‘Electroporation protocol - parameter screen’ section above with the following modifications for culturing and competent cell preparation.

<u>Culturing.</u> Overnight cultures were grown in 50-mL conical tubes covered with breathable membranes (ThermoFisher) instead of baffled flasks, and subsequently diluted into 10-30 mL of LB broth in a baffled flask to an OD_600_ of 0.08.

<u>Competent cell preparation.</u> Unconcentrated competent cells were prepared by washing 1 mL of culture three times with 670 μL of the appropriate cell suspension buffer and resuspending the final pellet in 1 mL of the same buffer. Centrifugation between wash steps was performed in sterile 1.5-mL Eppendorf tubes at 10,000 RCF for 2 minutes at room temperature (Eppendorf 5415D Centrifuge).

**Sanger sequencing**

Eton Bioscience was used to perform Sanger sequencing of single colonies from four-plasmid library transformants.

**Electroporation protocol - 10- and 17- plasmid library**

All steps were performed as described in the ‘Electroporation protocol - 24-condition screen’ section above using plasmid library for DNA input (**Supplementary Table 1**).

### NGS analysis of 17-plasmid library transformation

<u>Amplicon sequencing.</u> The 17-plasmid library hits were identified from the liquid selection plates as samples with ΔOD_600_ >0.25 (**Supplementary Figure 16**). Selected samples were pelleted (5 minutes, 5000 RCF at RT) stored at −20°C. gDNA extraction was performed using the MagMAX™ Viral/Pathogen Nucleic Acid Isolation Kit (Applied Biosystems). A control sample containing DNA but no cells, was processed to determine a minimum threshold of enrichment for positive hits.

All primers used can be found in **Supplementary Table 4**. The same plasmid barcode amplicon library constructs used for NGS screening of the POSSUM library of plasmid ORIs^43^ were constructed here, with some procedural modifications. All PCR reactions were done in a 20 μL volume using the KAPA HiFi HotStart PCR Kit (Roche) according to the manufacturer’s protocol: 4 μL KAPA HiFi Buffer (5X), 0.6 μL 10 mM KAPA dNTP Mix (10 mM), 0.6 μL Forward Primer (10 μM), 0.6 μL Reverse Primer (10 μM), 0.4 μL KAPA HiFi HotStart DNA Polymerase (1 U/μL), and the remaining 13.8 μL volume split between the appropriate template DNA volume and nuclease-free water. 1 μL of 5 ng/μL gDNA input was added to PCR 1; 1 μL was still added for any samples with less than 5 ng/μL starting concentration. An initial test qPCR was completed with these PCR 1 reactions by including 1 μL 20x EvaGreen Dye (Biotium) and cycling in a qTOWER^3^ G thermal cycler (Analytik Jena) using the following thermal protocol: 95°C for 3 minutes and 30 cycles of 98°C denaturation for 20 sec, 64°C annealing for 15 sec, 72°C extension for 15 sec. The amplification curves produced in the qPCR were evaluated for each sample to determine an appropriate number of cycles for the actual productive PCR reaction. Productive cycle numbers were called as the number of cycles it took for the sample to reach an Intensity [I] value between 50000 and 75000 and varied between 17 and 27 cycles. Productive PCR reactions were completed for each sample with their respective cycle numbers with the same amplification recipe as the initial qPCR except without the EvaGreen Dye.

Then, 1 μL of a 10-fold dilution of the PCR 1 product was added to PCR 2 as template and the reactions were cycled in a qTOWER^3^ G thermal cycler using the following thermal protocol: 95°C for 3 min; 12 cycles of 98°C denaturation for 20 sec, 70°C annealing for 15 sec, 72°C extension for 15 sec; and 72°C for 1 min. PCR 2 products were spotchecked for the correct sized product of 220 bp on a 2% EGel EX (Invitrogen) before pooling together by equal volume. The pooled libraries were then purified through a double-sided AMPure XP (Beckman Coulter) bead clean according to the manufacturer’s protocol using an initial 0.7x bead to sample volume ratio to exclude large fragments above 400 bp, followed by taking the supernatant and adding beads for a 1.2x final total bead to sample volume ratio to exclude small fragments below 150 bp. The concentration of the cleaned sample pool was measured using the Qubit 1X dsDNA High Sensitivity Kit (Invitrogen) and quantified with the KAPA Library Quantification Kit for Illumina Platforms (Roche). The rest of the library preparation was completed following the Denature and Dilute Libraries Guide for MiniSeq System by Illumina, then loaded and sequenced on an Illumina MiniSeq in a 2 x 75 bp paired-end run.

<u>Sequencing analysis.</u> The amplicon sequences from our transformant pools were analyzed in a similar manner as previously described for a high-throughput NGS screen using the POSSUM library of plasmid ORIs^43^. Briefly, the data was quality-trimmed and adapter-trimmed using BBDuk^74^, and the respective barcodes of each ORI were matched using exact matches only with VSEARCH^75^, as implemented in the barcode_quantification.py script on the GitHub repository associated with the POSSUM toolkit: <u>github.com/cultivarium/ORI-marker-screen</u>. A DNA-only control was sequenced, along with triplicate negative controls (wells with no transformant growth). The abundance of each ORI in the plasmid pools was normalized to the nonfunctional dummy ORI in the negative controls. A cutoff of five-fold the maximum ratio of any ORI to the dummy observed across the negative controls was used as previously described^43^, and for this experiment this cutoff was an enrichment of at least 9.2x over the dummy ORI. Amplicon data from transformant pools was then analyzed in the same manner, and all ORIs more than 9.2-fold more abundant than the dummy ORI were called as present in the sample.

### Active learning algorithm development

The active learning process was guided by a Bayesian optimization algorithm for electroparameter selection, implemented using the optuna package^76^ in Python. The chosen parameter space for waveform, voltage, resistance, capacitance, pulse duration, buffer, and buffer concentration are summarized in **Supplementary Table 5**. Each cycle of optimization consisted of running two 96-well plates simultaneously. The initial plates were designed with a Quasi Monte Carlo sampler with the QMCSampler class, an algorithm which optimizes for even sampling of the parameter space before any data has been collected for that particular strain. The subsequent optimized plates were designed with a Tree-structured Parzen Estimator^77^, an estimator that fits a Gaussian Mixture Model (GMM) to the parameters associated with positive transformation efficiencies in prior data, and another GMM to the parameters associated with negative transformation efficiencies, and then suggests new parameter values that maximize the difference between the two models. The TPESampler was implemented using the TPESampler class in the optuna package with the parameter constant_liar=True to facilitate batch suggestions for the cycle without converging on a single suggestion. All data collected from previous plates for each cycle were added as individual trials to the TPESampler study. Each optimization cycle is therefore composed of two 96-well plates, modeled as an TPESampler study with all prior data provided to the algorithm as individual trials, and then suggestions for new parameters are generated for both plates in that cycle.

Suggestions for electroporation parameters were generated in batch format. The sampler suggests parameters for diagonal wells representing a new row and column value, and then iteratively adding all corresponding wells as individual trials to the study object by filling the associated rows and columns for that row, moving orthogonally across the plate. Parameters for the cell-only negative control column (Column 12) were kept constant in all electroplates: Exponential decay, 1000 V, 25 µF, 200 Ω. Optimization was performed with the goal of maximizing TE, and all TE results from prior plates informed the sampler during generation of the subsequent plates. For the third cycle of optimization, the plate design was modified so that parameters resulting in the highest TE from the previous plate were included on the resulting plate. The plot_param_importances function in optuna was used to calculate functional ANOVA hyperparameter importances^69^. Cycle electroplate parameters were exported in JSON format for machine interpretation. Code for this implementation is made available at <u>github.com/cultivarium/electroporation-bayesian-optimization</u>.

### Active learning electroporation

Experiments were performed using a custom-fabricated robotic platform which integrates the custom-built electroporator. Unless otherwise stated, all protocol steps were performed at room temperature and on the robotic platform.

<u>Culturing</u>. Initial *C. necator* cultures were inoculated manually directly from glycerol stock into 50 mL of NB media and incubated at 30°C shaking for 16-24 hours until an OD_600_ of ∼0.7 was reached. Then, 1 mL of culture was aliquoted into each well of 4 columns of a 96-deep well plate (32 wells total) and placed on the robotic system deck for downstream competent cell preparation, electroporation, and recovery.

<u>Buffer plate preparation</u>. Buffer preparation was performed manually as described in the “**Electroporation buffer preparation”** section above. Into two 8-channel reservoirs, 15-mL of the 16 buffers selected by the Bayesian optimizer was aliquoted into the appropriate channel. The buffer plates were placed on the robotic platform for competent cell preparation.

<u>DNA preparation.</u> On ice, 900 µL of 100 ng/µL pGL2_270 plasmid DNA was thawed and transferred to a cooling block on the robotic platform. In two 0.2-cm electroplates, 5 μL of plasmid DNA was added to each well of columns 1-11.

<u>Competent cell preparation.</u> Cells were pelleted by centrifugation at 4200 RCF for 5 minutes (Sorvall ST-1 R with H-Flex Rotor). Then, ∼950 μL of supernatant was removed. Cells were washed with 700 µL of buffer in the corresponding row. Centrifugation and wash steps were repeated for a total of four washes, but in subsequent wash steps ∼700 μL of supernatant was removed. This yielded ∼750 μL of competent cells per well, sufficient for 7 transformation reactions.

<u>Electroporation and recovery.</u> Row-wise, 95 μL of competent cells were aliquoted into each well of column 1-12 of the two 0.2-cm electroplates. Column 12 contained no DNA and thus acted as a negative control. Electroporation parameters for each column and plate as suggested by the Bayesian optimizer were programmed into the robotics user interface. Following electroporation, 100 µL was immediately transferred from the electroplate into the recovery plate containing 900 µL of NB media in every well. The recovery plate was sealed with a breathable membrane and incubated at 30°C shaking for 20 hours.

<u>Selection</u>. After 20 hours of recovery, both agar and liquid selection was performed. On the robotic platform, cells were diluted 10-fold up to 10^-2^ in Cycle 1 and up to 10^-3^ in Cycle 2 and 3 in 900 µL of PBS. Then, 50 µL from each well of the three dilution plates (10^0^-10-^2^ in Cycle 1 and 10^-1^-10^-3^ in Cycle 2 and 3) was transferred to two 96-deep well plates containing either 950 µL of NB media supplemented with 50 µg/mL kanamycin (selective) or 950 µL of NB media (non-selective). From the same dilution plates, 5 µL was spot plated onto non-selective and selective solid media using the Nayo N96. All plates were incubated at 30°C. Liquid plates were grown with shaking for 20 hours while solid plates were incubated until colonies appeared (1-2 days).

<u>Data collection</u>. Colonies on solid plates were counted manually. OD_600_ of liquid cultures were measured using 100 µL of sample on the Epoch 2 plate reader on the robotic platform.

### Comparison of active learning protocols to literature

Experiments were performed *manually* using electroporation cuvettes.

<u>Culturing</u>. Initial *C. necator* culture was inoculated directly from glycerol stocks into 50 mL of SOB media and incubated at 30°C shaking for 16 hours. Saturated overnight culture (OD_600_ 4.95) was diluted into three flasks containing 50 mL of SOB to OD_600_ 0.1 and incubated until an OD_600_ of 0.6 was reached.

<u>Competent cell preparation.</u> Flasks were incubated on ice for 10 minutes. Cultures were transferred to 50-mL conical tubes and centrifuged at 5000 x *g* at 4°C for 10 minutes. Supernatant was removed, and cells were washed according to the corresponding protocol below:

*Cycles 1 and 3*. Cells were resuspended in 50 mL of Milli-Q water by pipetting up and down with a 50-mL serological pipette. The centrifugation and washing steps were repeated for a total of three washes. The cells were pelleted by centrifugation as before. The cell pellet was finally resuspended in 500 µL for 100-fold concentration, and 4x 100 µL aliquots were transferred to eppendorf tubes (100x cells). The remaining volume was brought up to 10 mL and 4x 100 µL aliquots were transferred to eppendorf tubes (1x cells). Then, 5 µL of DNA (pGL2_270) was added to each eppendorf tube (except 1 aliquot for the negative control where molecular grade water was added instead). Tubes were flicked to mix, and the contents were transferred to 0.2-cm cuvettes and incubated on ice for 5 minutes.

*Literature.* Cells were prepared as previously described^68^. Cells were resuspended in 25 mL of 50 mM CaCl_2_ by briefly using a vortex and incubated for 15 minutes on ice. The cells were pelleted by centrifugation as before. Cells were washed twice using 25 mL and 15 mL of ice-cold 0.2 M sucrose, respectively. The cell pellet was finally resuspended in 500 µL for 100-fold concentration, and 4x 100 µL aliquots were transferred to eppendorf tubes (100x cells). The remaining volume was brought up to 10 mL and 4x 100 µL aliquots were transferred to eppendorf tubes (1x cells). Then, 5 µL of DNA (pGL2_270) was added to each eppendorf tube (except 1 aliquot for the negative control where molecular grade water was added instead). Tubes were flicked to mix, and the contents were transferred to 0.1-cm cuvettes and incubated on ice for 5 minutes.

<u>Electroporation and recovery.</u> Cells were electroporated with the following parameter settings: Cycle 1 #1 - EX, 2.715 kV, 25 μF, 550 Ω; Cycle 3 #1 - EX, 2.555 kV, 25 μF, 175 Ω; Cycle 3 #2 - EX, 1.750 kV, 25 μF, 175 Ω; Literature - EX, 1.150 kV, 25 μF, 200 Ω. Following electroporation of each sample, 900 µL of SOB media was immediately added to the cuvette. Cells were transferred to a 2 mL centrifuge tube for recovery with shaking at 30 °C for 2 h.

<u>Selection</u>. Transformants were selected on solid media supplemented with KAN (50 µg/mL), plating 100 µL of 10^0^ and 10^-1^ (for 1X cells) or 10^-2^ and 10^-3^ (for 100X cells) on full Petri dishes, as well as spot plating 5 µL up to 10^-8^ dilution. Plates were incubated for 20 hours and colonies were counted manually.

## Data and Resource Availability

The POSSUM toolkit is available via Addgene (<u>POSSUM Toolkit</u>, Kit ID #1000000234) and sequencing analysis scripts for the POSSUM ORI screens are available at <u>github.com/cultivarium/ORI-marker-screen/</u>. Code for Bayesian optimizer is available at <u>github.com/cultivarium/electroporation-bayesian-optimization</u>.

## Author Contributions

A.D.C., S.L.B., N.O. and H.H.L. designed the project. S.H., A.L., and A.D.C. performed and analyzed initial parameter selection experiments and contributed to an early draft of the manuscript. A.L. and S.L.B. performed and analyzed library-scale experiments. S.L.B. performed active learning experiments. A.C.C. developed the electroporation Bayesian optimizer. S.L.B. and A.C.C. performed analysis of active learning data. M.N. and A.L. automated electroporation procedures. A.E. performed DNA extractions. J.L. performed DNA sequencing workflows. A.D.C., L.L., and M.N. oversaw construction of the custom instrumentation. S.L.B., A.D.C., C.G., H.H.L., and N.O. performed data analysis. A.C.C. analyzed library delivery data. S.L.B., N.O., and H.H.L. wrote the manuscript. All authors have read and approved the manuscript.

## Acknowledgements

We thank all members of the Cultivarium team for discussions throughout this project. Cultivarium acknowledges support from Eric and Wendy Schmidt as a Convergent Research Focused Research Organization (FRO).

