## Supplementary Figures and Tables for "Active learning guides automated discovery of DNA delivery via electroporation for non-model microbes"

**Supplementary Figures** **2**
Supplementary Figure 1. Plasmid map of pAKgfp1-kan. 2
Supplementary Figure 2. Transformation of *E. coli* using 0.2-cm single cuvette or 0.2-cm 96-well electroplate. 3
Supplementary Figure 3. Effect of wash buffer on transformation efficiency. 3
Supplementary Figure 4. Effect of voltage on cell survival. 4
Supplementary Figure 5. Effect of resistance and capacitance on transformation efficiency. 4
Supplementary Figure 6. Effect of waveform on transformation efficiency. 5
Supplementary Figure 7. Effect of post-electroporation recovery on transformation efficiency in *E. coli*. 6
Supplementary Figure 8. Effect of competent cell concentration on transformation efficiency. 7
Supplementary Figure 9. 24-condition electroporation screen on seven bacteria using a custom device. 8
Supplementary Figure 10. Solid agar selection of 24-condition electroporation screen. 9
Supplementary Figure 11. Liquid selection of 24-condition electroporation screen. 10
Supplementary Figure 12. Optimization and validation of electroporation protocols in *H. elongata*. 11
Supplementary Figure 13. Reproducibility of electroporation screening across waveforms and biological replicates. 12
Supplementary Figure 14. Liquid selection for 24-condition electroporation screen using POSSUM plasmid library delivery to five bacteria. 13
Supplementary Figure 15. Solid selection for 24-condition electroporation screen using POSSUM plasmid library delivery to five bacteria. 14
Supplementary Figure 16. *P. alcaliphila* plasmid library ORI sequencing landscape. 15
Supplementary Figure 17. Correlation between solid and liquid selection for active learning. 16
Supplementary Figure 18. Buffer, waveform, and voltage conditions tested during three cycles of active learning. 16
**Supplementary Tables** **17**
Supplementary Table 1. Plasmids and plasmid libraries used in this study. 17
Supplementary Table 2. Strains used in this study. 18
Supplementary Table 3. Previously reported electroporation conditions for strains used in this study. 18
Supplementary Table 4. Primers used in this study. 19
Supplementary Table 5. Electroporation parameter constraints for Bayesian optimization algorithm. 19
Supplementary Table 6. Growth conditions and selective antibiotic concentrations used for bacteria in this study. 20
Supplementary Table 7. POSSUM toolkit parts for construction of chloramphenicol 10-plasmid pool. 20
Supplementary Table 8. Experimental overview of 24-condition screen. 21

#### Supplementary Figures

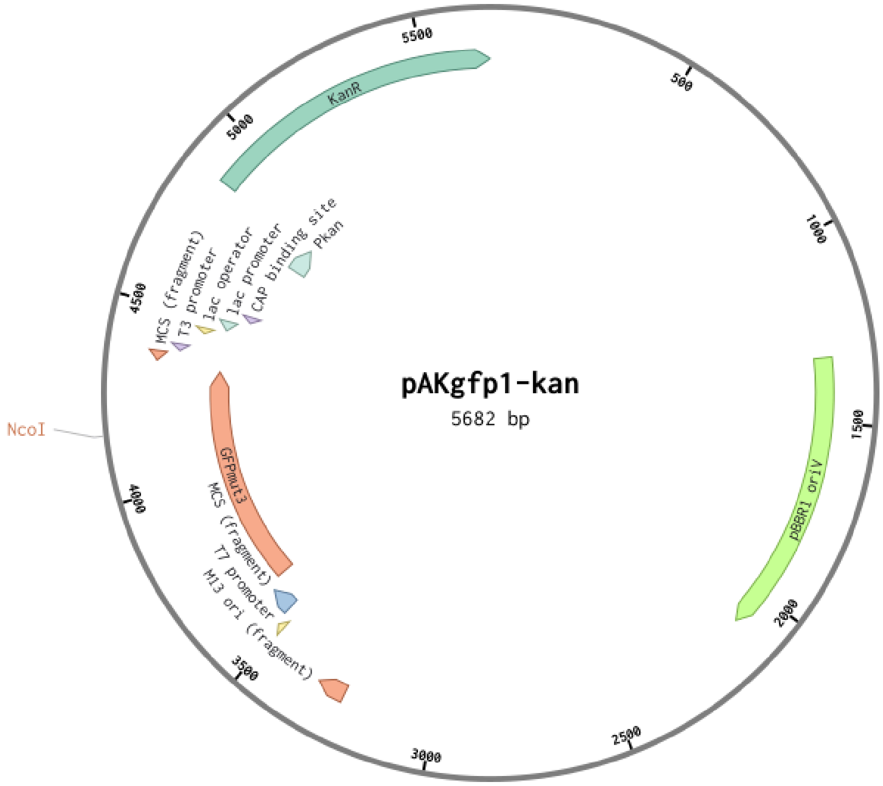


**Supplementary Figure 1. Plasmid map of pAKgfp1-kan.** Main plasmid components include a kanamycin antibiotic resistance gene (KanR), the pBBR1 broad-host-range origin of replication (pBBR1 oriV), and a GFP cassette (GFPmut3) under the T7 promoter.

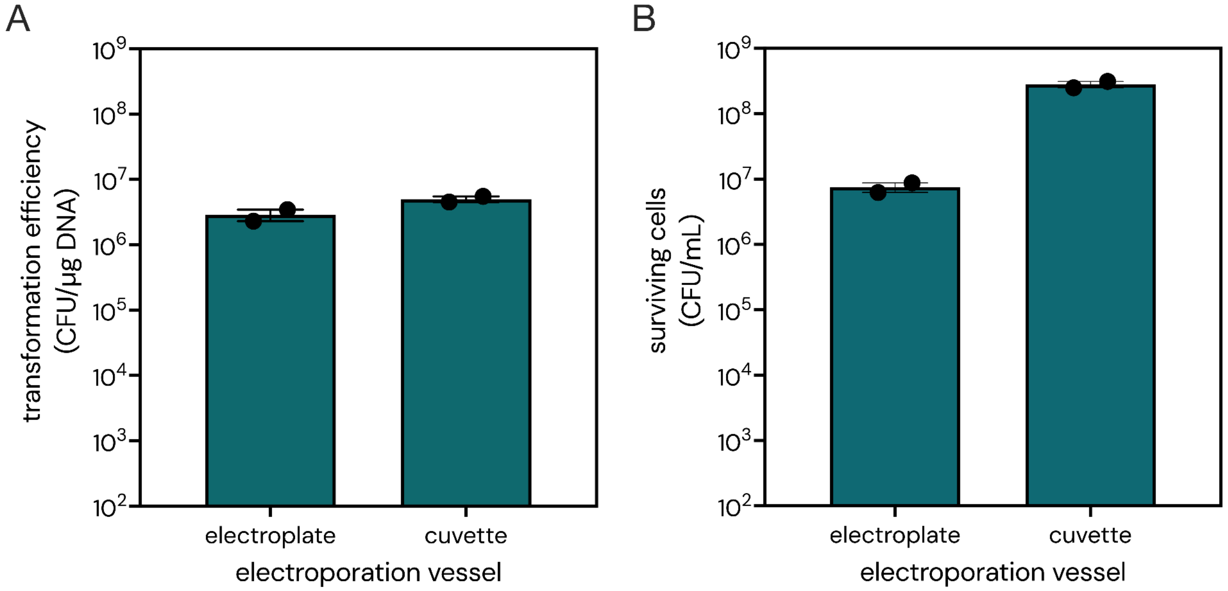


**Supplementary Figure 2. Transformation of *E. coli* using 0.2-cm gap cuvette or 0.2-cm 96-well electroplate.** (**A**) Transformation efficiency. (**B**) Total surviving cells. Competent cells were concentrated 100-fold of the original culture volume and samples were electroporated at 3 kV, 200 Ω, and 25 µF. After electroporation, 1 mL of recovery media was added immediately and cells were recovered for 1 hour. Transformants were selected on kanamycin plates to determine transformation efficiency. Data are the average of two biological replicates. Error bars represent standard error.

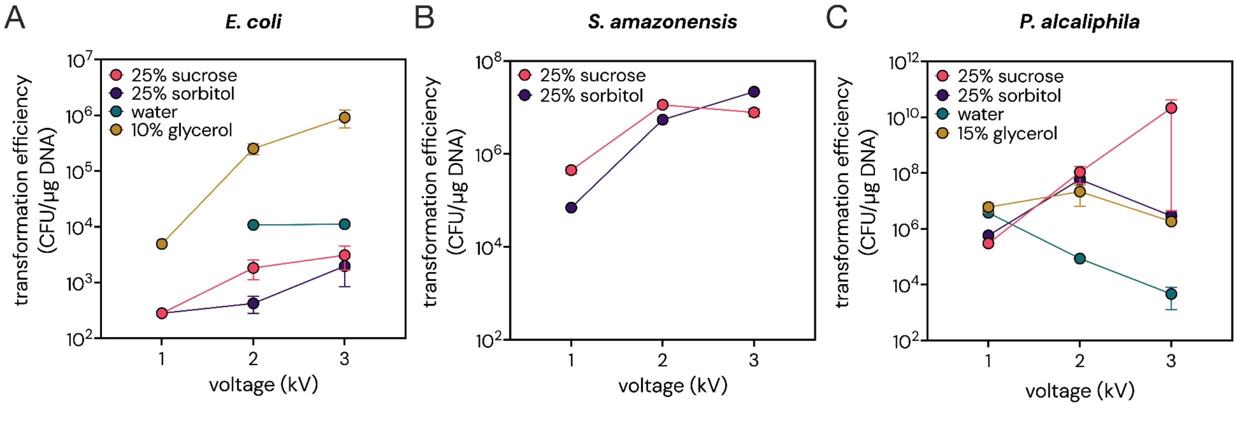


**Supplementary Figure 3. Effect of wash buffer on transformation efficiency.** Transformation efficiency of pAKgfp1 in (**A**) *E. coli* and pAKgfp1-kan in (**B**) *S. amazonensis* and (**C**) *P. alcaliphila*. Competent cells were concentrated 100-fold (*E. coli*, *P. alcaliphila*) or 60-fold (*S. amazonensis*) the original culture volume (50-100 mL). Samples were electroporated using exponential waveform at 200 Ω and 25 µF using the BTX Gemini x2 Electroporator in 96-well electroplates. After electroporation, *E. coli*, *S. amazonensis,* and *P. alcaliphila* cells were recovered for 1 hour, 2 hours, and 1.5 hours, respectively and plated on ampicillin (*E. coli*) or kanamycin (*S. amazonensis, P. alcaliphila*). Data are the average of two biological replicates. Error bars represent standard error.

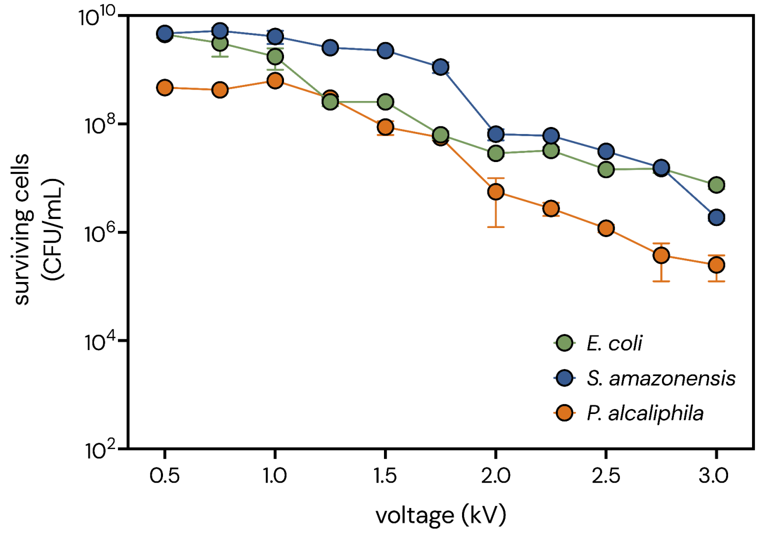


**Supplementary Figure 4. Effect of voltage on cell survival.** Number of surviving cells following transformation of pAKgfp1-kan into *E. coli*, *S. amazonensis,* and *P. alcaliphila*. Competent cells were concentrated 100-fold (*E. coli* and *P. alcaliphila*) or 60-fold (*S. amazonensis*) of the original culture volume. Buffers used for *E. coli, S. amazonensis* and *P. alcaliphila* were 10% glycerol, 25% sorbitol, and 25% sucrose, respectively. After electroporation, cells were recovered for 1 h, 2 h and 1.5 h for *E. coli, S. amazonensis* and *P. alcaliphila* in LB, respectively. All samples were electroporated using exponential decay waveform at 200 Ω and 25 μF. Data are the average of two biological replicates. Error bars represent standard error.

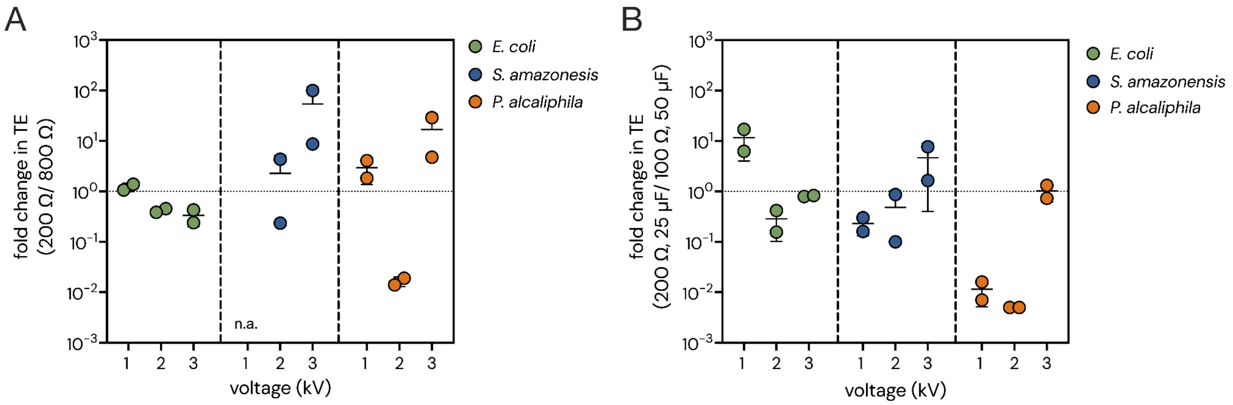


**Supplementary Figure 5. Effect of resistance and capacitance on transformation efficiency.** The exponential decay rate can be tuned by altering the resistance (R) and capacitance (C), which is expressed as the time constant (τ = RC). (**A**) Fold change in transformation efficiency (TE) between low (200 Ω) and high (800 Ω) R with a constant C (25 μF) across three voltages (1, 2, and 3 kV). (**B**) Fold change in TE using two different C and R settings both producing a time constant (τ) of ~5 ms: 200 Ω and 25 μF or 100 Ω and 50 μF. (**A-B**) In all experiments, *E. coli* was washed with 10% glycerol and concentrated 100-fold, *S. amazonensis* cells were washed with 15% sorbitol and concentrated 60-fold, and *P. alcaliphila* cells were washed with 15% glycerol and concentrated 100-fold. Data are the average of two biological replicates. Error bars represent standard deviation.

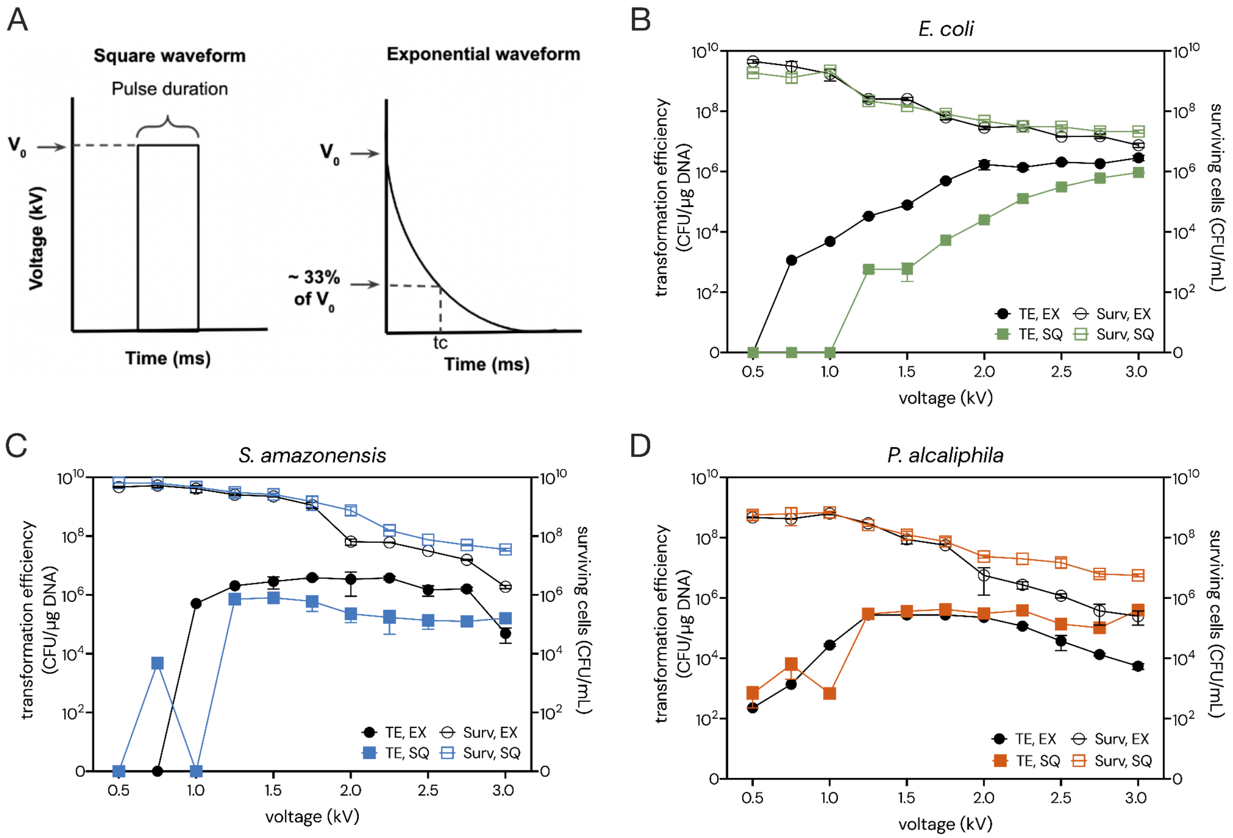


**Supplementary Figure 6. Effect of waveform on transformation efficiency.** (**A**) Representation of square wave and exponential decay wave pulses. (**B-D**) Transformation efficiency and surviving cells for (**B**) *E. coli*, (**C**) *S. amazonensis*, and (**D**) *P. alcaliphila*. In each plot, colored squares represent square waveform (SQ) and black circles represent exponential decay (EX). Open shapes represent surviving cells (Surv, right Y-axis) and filled shapes represent transformation efficiency (TE, left Y-axis). Cells were concentrated 100x (*E. coli* and *P. alcaliphila*) or 60x (*S. amazonensis*) the original culture volume. Resistance and capacitance were kept constant at 200 Ω and 25 µF for all exponential decay experiments. Pulse duration was set to 0.6 ms in all square wave experiments. Buffers used for *E. coli, S. amazonensis* and *P. alcaliphila* were 10% glycerol, 25% sorbitol, and 25% sucrose, respectively. Data are the average of two biological replicates. Error bars represent standard error.

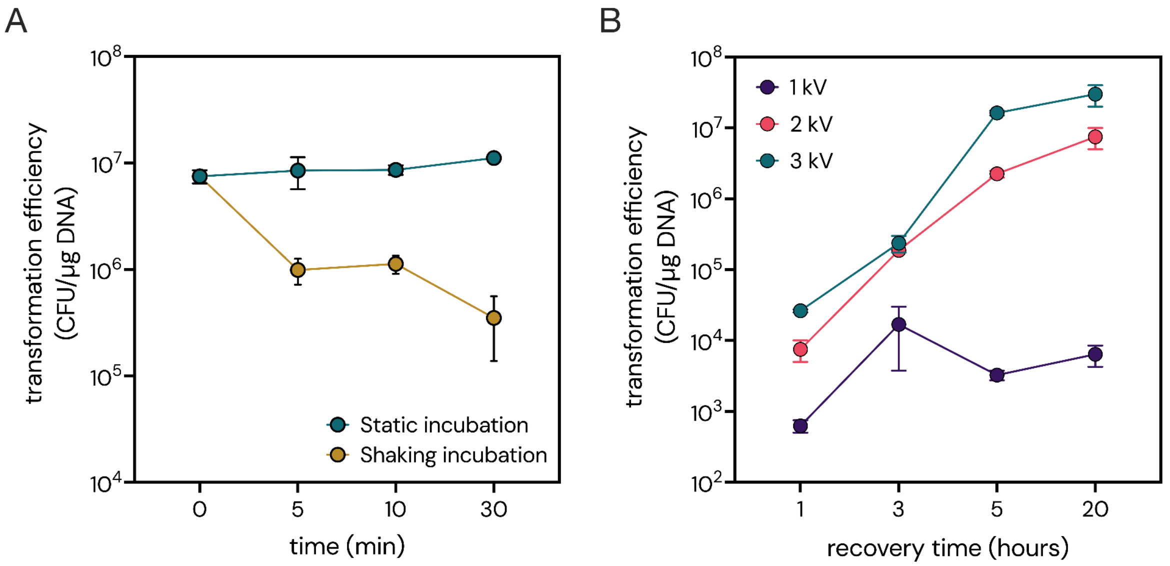


**Supplementary Figure 7. Effect of post-electroporation recovery on transformation efficiency in *E. coli*.** (**A**) Effect of delay in recovery start time on transformation efficiency. Transformed cells either 1) had recovery media added immediately following electroporation and were incubated stationary at room temperature until the indicated time when they were transferred to the incubator (static incubation) or 2) did not have recovery media added until the indicated time when they were transferred to the incubator (shaking incubation). Once recovery media was added (1 mL of LB), all cells were recovered for 1 hour shaking at 225 RPM at 37°C. (**B**) Transformation efficiency achieved with 1-20 hour recovery times at three voltages (1, 2, and 3 kV). (**A-B**) Competent cells were washed with 10% glycerol, concentrated 100-fold of the original culture volume, and electroporated at resistance and capacitance of 200 Ω and 25 µF. Data are the average of two biological replicates. Error bars indicate standard error.

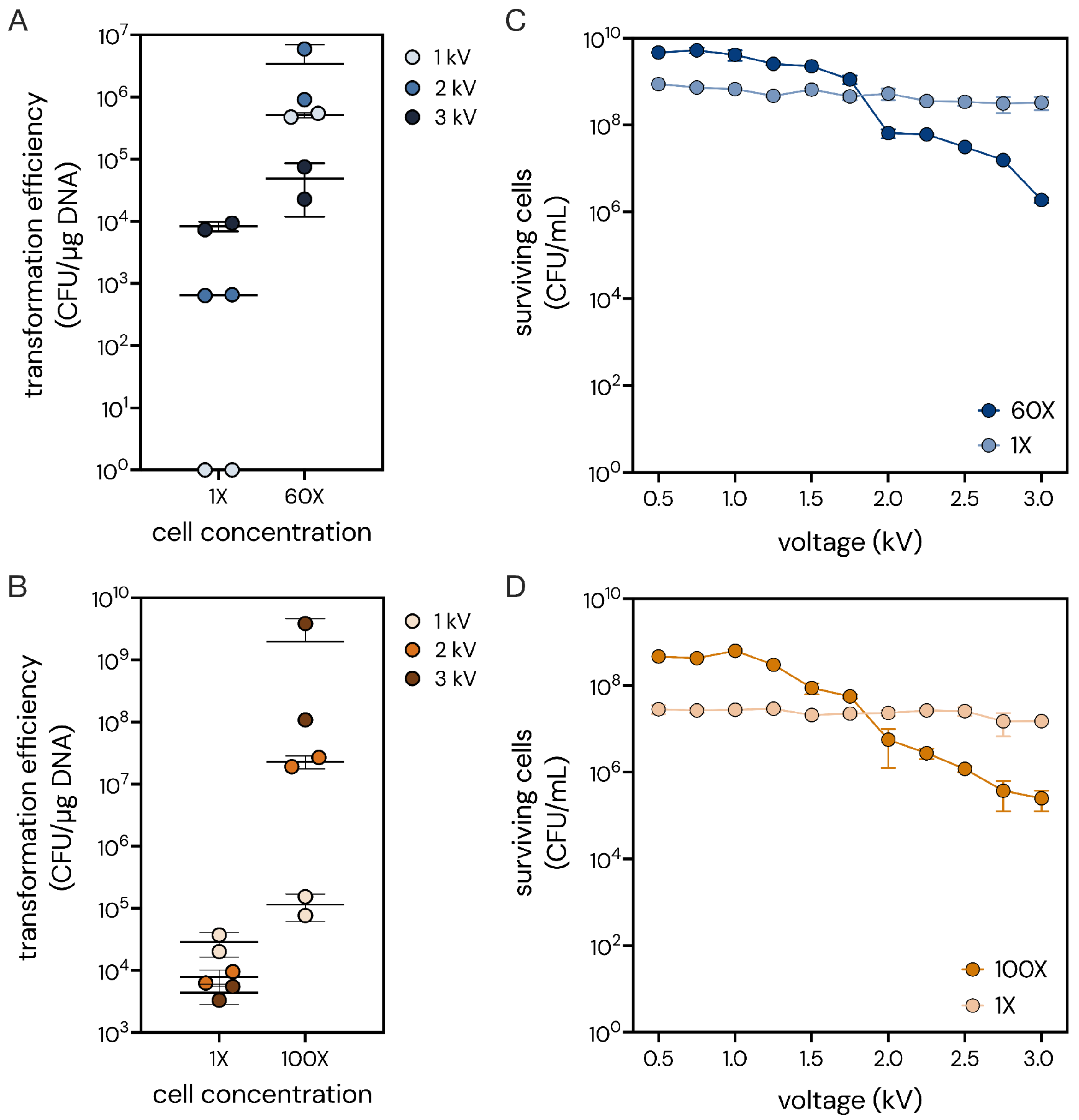


**Supplementary Figure 8. Effect of competent cell concentration on transformation efficiency.** (**A-B**) Transformation efficiency of unconcentrated (1X) versus concentrated (60X or 100X) cells for (**A**) *S. amazonensis* and (**B**) *P. alcaliphila* at 1, 2 and 3 kV. **(C-D)** Survival of unconcentrated (1X) versus concentrated (60X or 100X) cells following electroporation at a range of voltages in (**C**) *S. amazonensis* and (**D**) *P. alcaliphila*. Concentrated and unconcentrated cells were grown in 50-100 mL or 10-30 mL cultures, respectively, to $OD_{600}$ of ~1.0, washed three times with the selected buffer (*S. amazonensis* using 25% sorbitol, *P. alcaliphila* using 25% sucrose), and resuspended with buffer to either the original volume (1X) or to a smaller volume (60X or 100X). All experiments were performed using exponential waveform with constant resistance and capacitance settings (200 Ω and 25 µF, respectively). Cells were recovered for 2 hours prior to plating on selective media. Data are the average of two biological replicates. Error bars represent standard error.

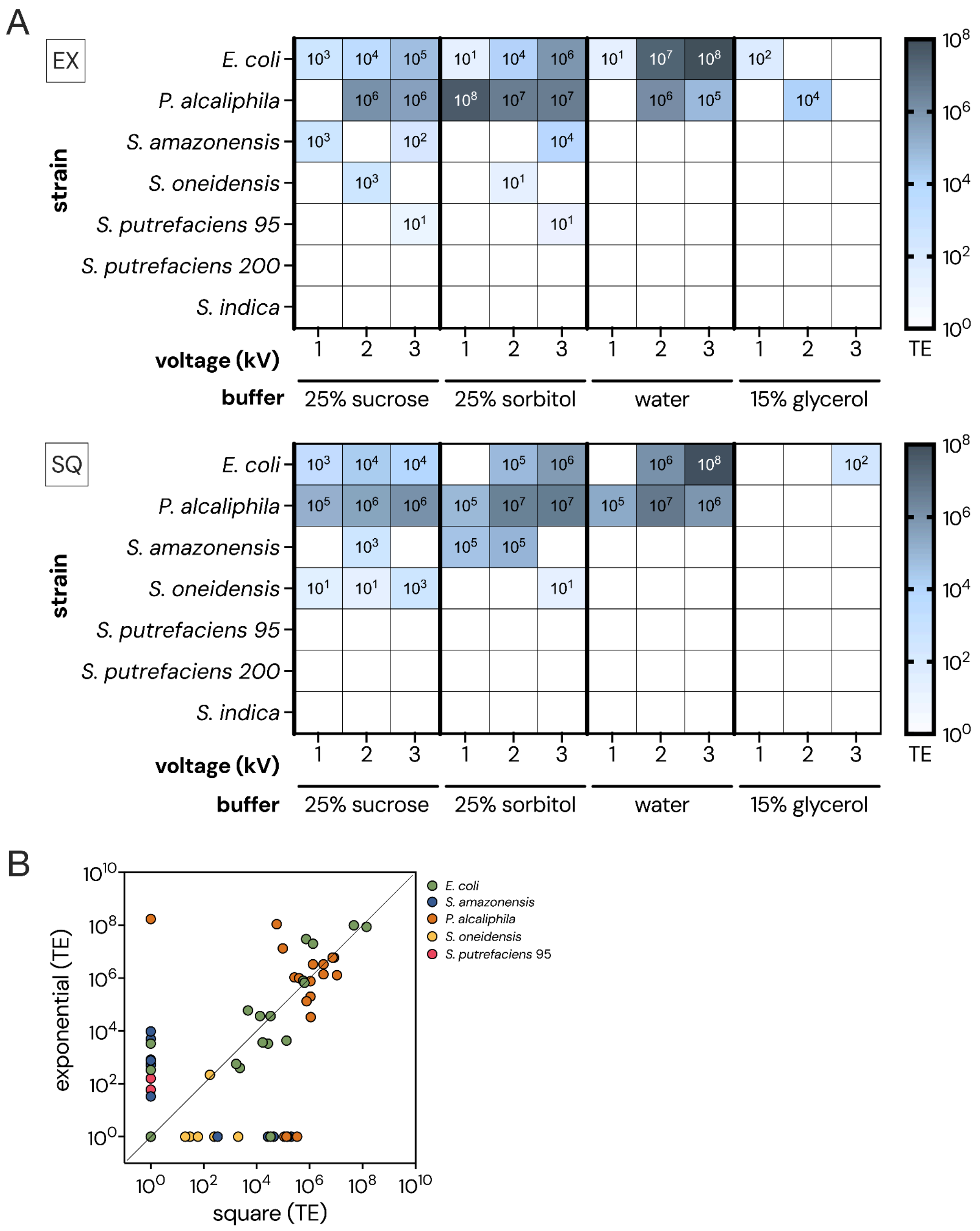


**Supplementary Figure 9. 24-condition electroporation screen on seven bacteria using a custom device.** (**A**) Transformation efficiency (TE) of 24-condition electroporation screen. Results are provided for both exponential decay (EX, top) and square (SQ, bottom) waveforms. (**B**) Correlation between TE using exponential decay and square waveforms for all bacteria tested in panel A (12 conditions were tested in each waveform). All screens were performed using low cell input protocol (unconcentrated cells) at constant resistance and capacitance (200 Ω and 25 µF, respectively). TE, transformation efficiency reported as CFU/µg DNA.

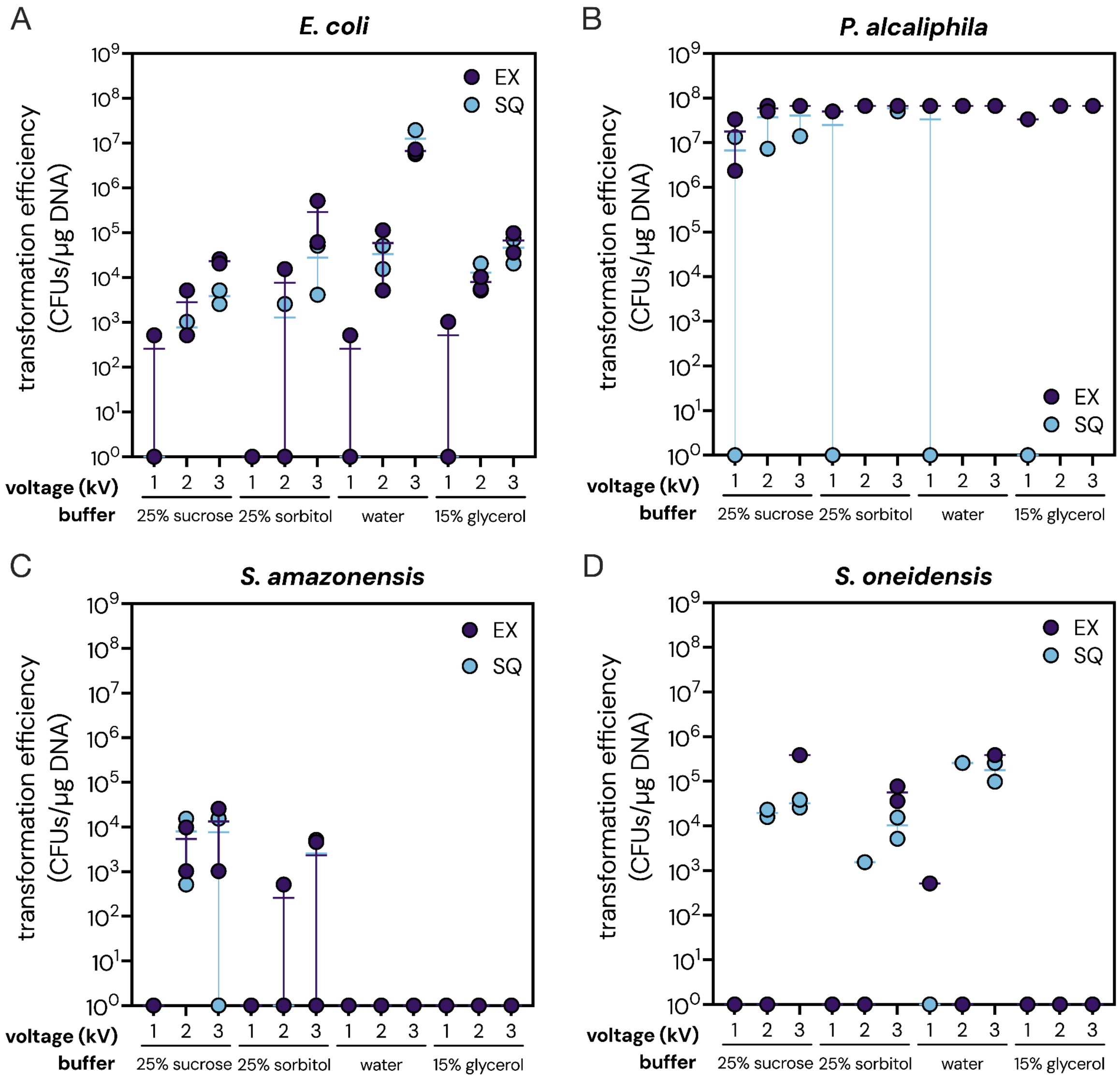


**Supplementary Figure 10. Solid agar selection of 24-condition electroporation screen.** (**A-D**) Transformation efficiency reported as CFU/µg DNA following electroporation screen of a single plasmid in (**A**) *E. coli*, (**B**) *P. alcaliphila*, (**C**) *S. amazonensis*, and (**D**) *S. oneidensis*. EX, exponential decay; SQ, square waveform. Data are the average of two biological replicates. Error bars represent standard deviation. All screens were performed using unconcentrated cells at constant resistance and capacitance (200 Ω and 25 µF, respectively).

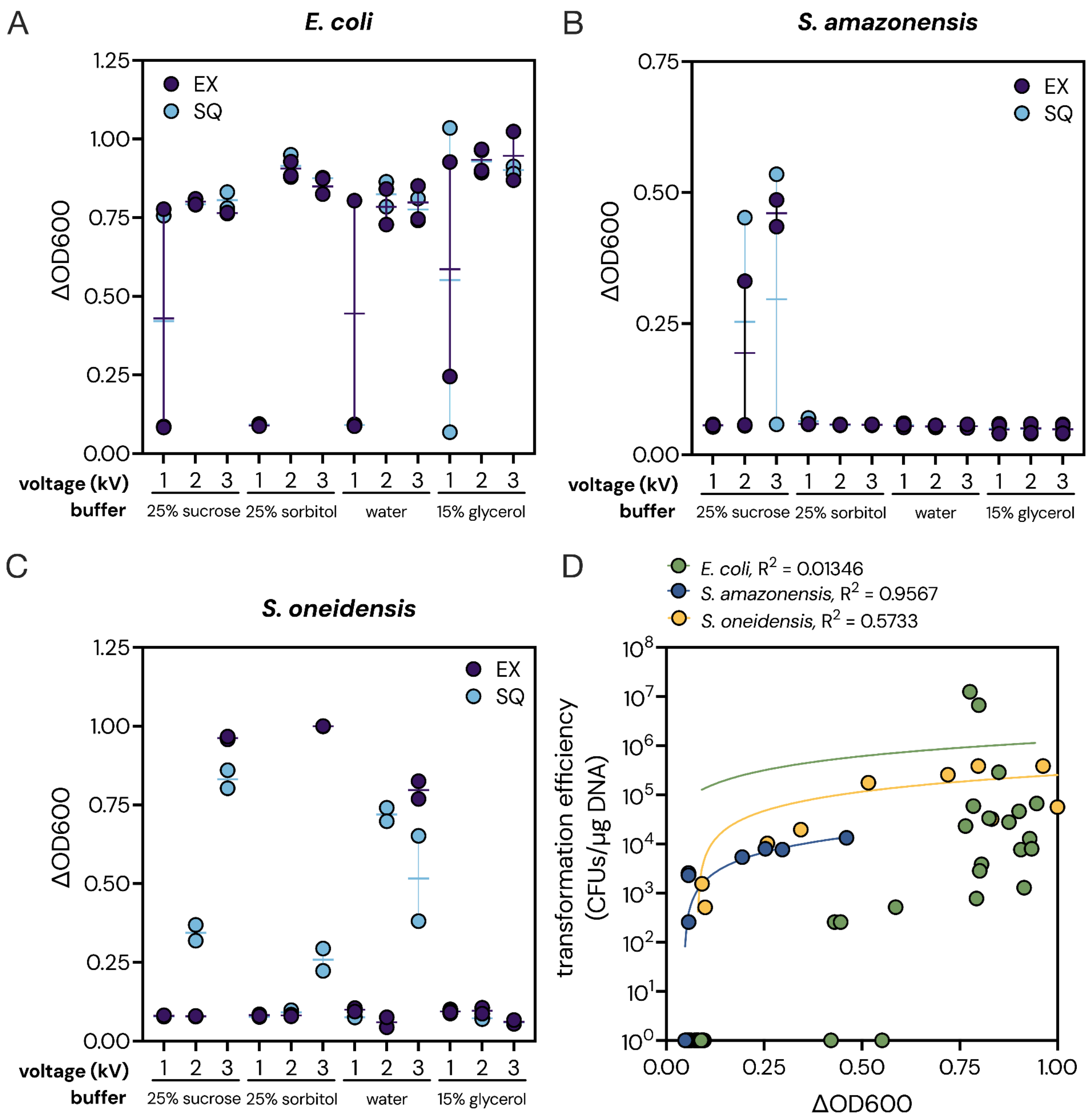


**Supplementary Figure 11. Liquid selection of 24-condition electroporation screen.** (**A-C**) Liquid selection results following electroporation screen using a single plasmid in (**A**) *E. coli*, (**B**) *S. amazonensis*, and (**C**) *S. oneidensis*. $\Delta OD_{600}$ is reported as the difference in $OD_{600}$ of the sample compared to a control containing electroporated cells but no DNA, both grown in selective media. (**D**) Correlation between liquid ($\Delta OD_{600}$) and solid (CFU/µg DNA) selection. Simple linear regression is shown for each strain. All screens were performed using unconcentrated cells at constant resistance and capacitance (200 Ω and 25 µF, respectively). EX, exponential decay; SQ, square waveform. Data are the average of two biological replicates. Error bars represent standard deviation.

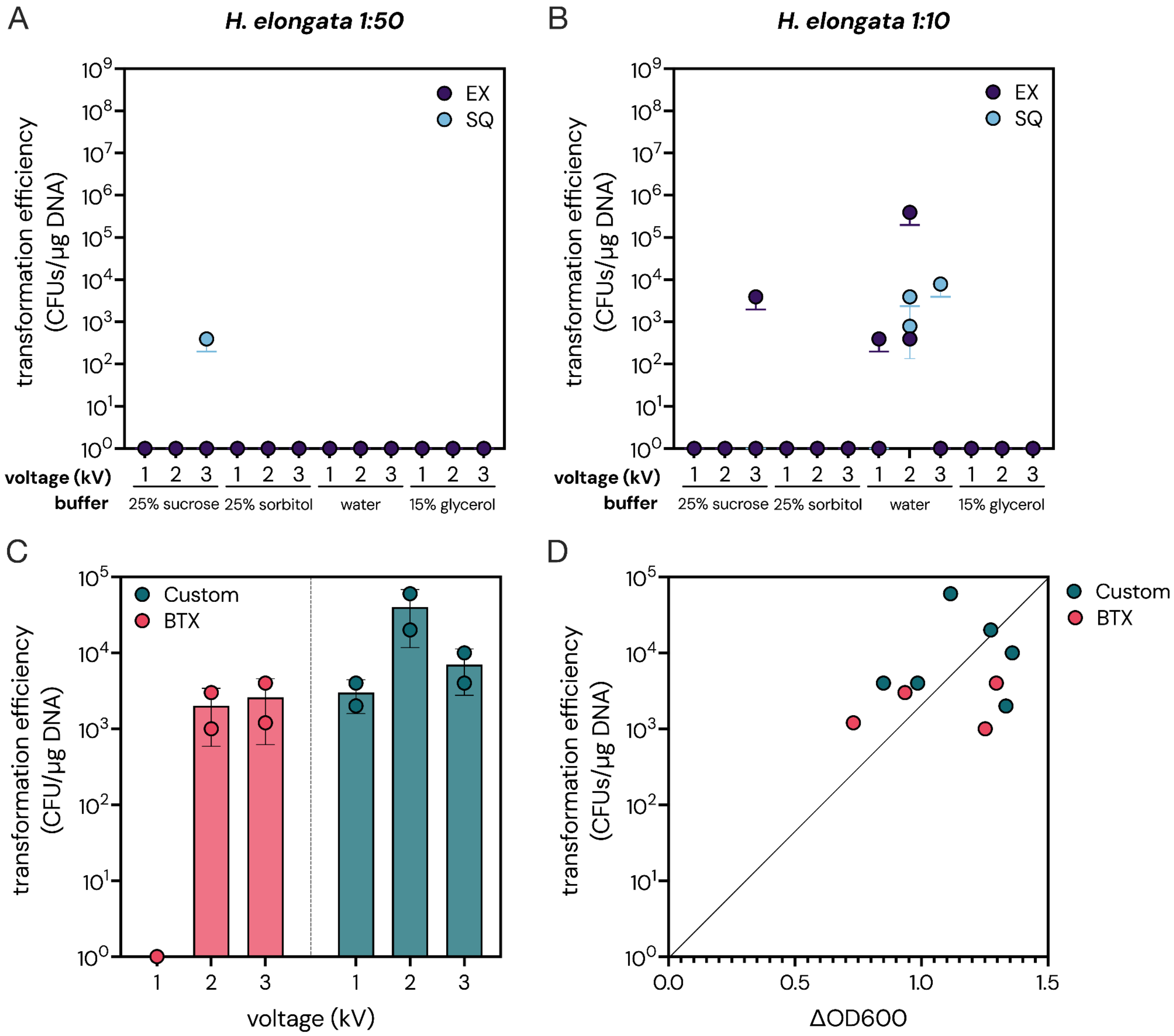


**Supplementary Figure 12. Optimization and validation of electroporation protocols in *H. elongata*.** (**A-B**) Transformation efficiency for *H. elongata* at varying cell densities: (**A**) using a standard protocol, where overnight culture was subcultured 1:50; or (**B**) using a modified protocol where overnight culture was subcultured 1:10. EX, exponential decay; SQ, square waveform. Data are the average of two biological replicates. Error bars represent standard deviation. (**C-D**) Single cuvette validation of *H. elongata* electroporation conditions. Cells were washed in Milli-Q and plasmid pGL2_168 (Addgene, 199100) was electroporated using square waveform in either single cuvettes (BTX) or our 96-well plate custom device (Custom) at the indicated voltages. (**C**) Transformation efficiency. (**D**) Correlation between transformation efficiency as measured on agar plates (Y-axis) and $\Delta OD_{600}$ as measured in liquid medium (X-axis). Data are the average of two biological replicates. Error bars represent standard deviation.

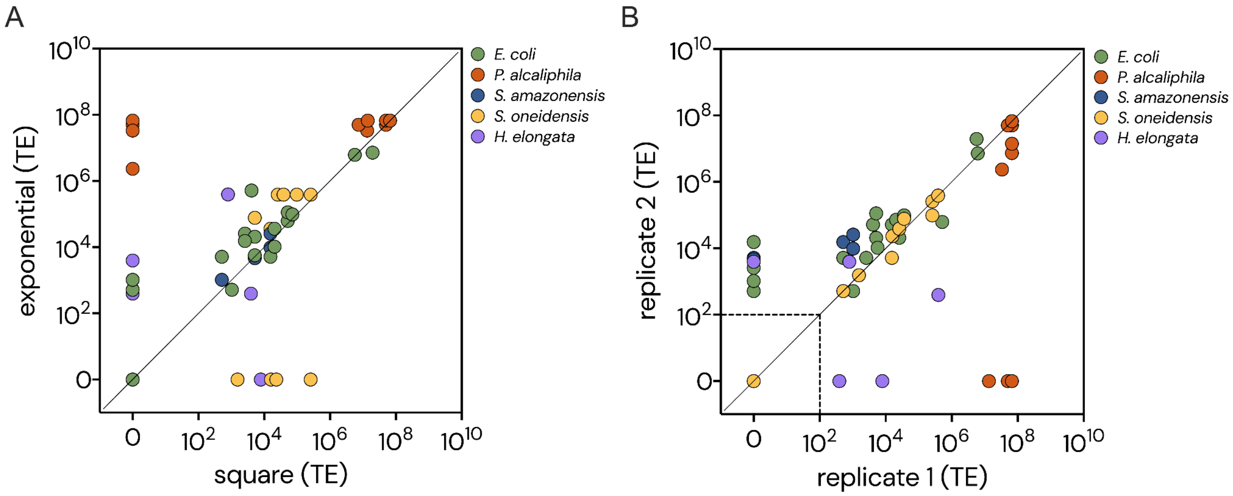


**Supplementary Figure 13. Reproducibility of electroporation screening across waveforms and biological replicates. A**) Correlation of transformation efficiency using exponential decay or square waveforms. (**B**) Comparison of transformation efficiency between two biological replicates of the 24-condition electroporation screen (including both exponential decay and square waveforms) from agar selection. The limit of detection for transformants is indicated by a dashed line. TE, transformation efficiency reported as CFU/µg DNA.

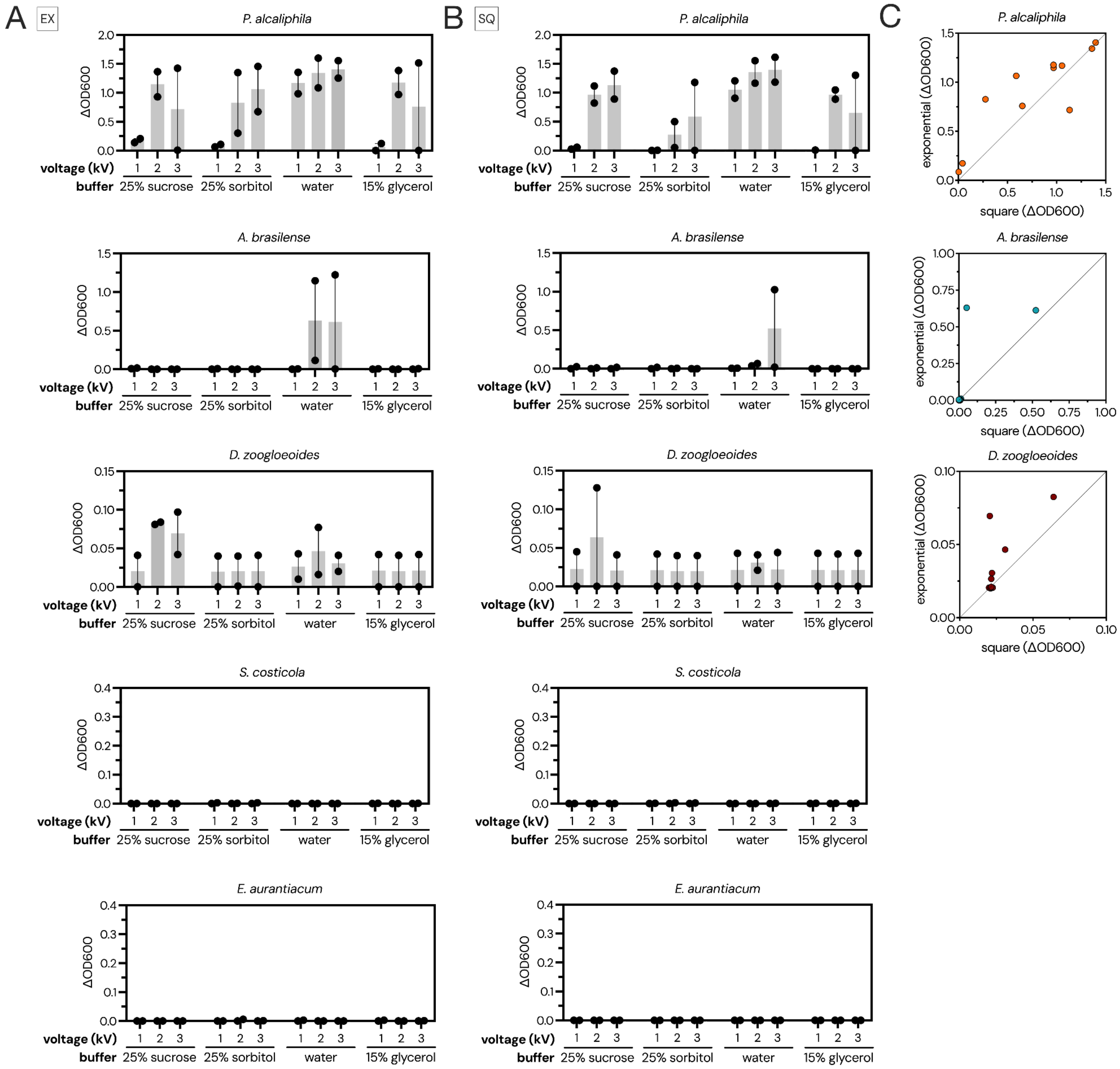


**Supplementary Figure 14. Liquid selection for 24-condition electroporation screen using POSSUM plasmid library delivery to five bacteria.** Results of liquid selection following plasmid library delivery to *P. alcaliphila, A. brasilense, D. zoogloeoides, S. costicola,* and *E. aurantiacum.* $\Delta OD_{600}$ represents the difference in $OD_{600}$ between samples with or without DNA (control). Data are the average of two biological replicates. (**A-B**) $\Delta OD_{600}$ using (**A**) exponential decay waveform (EX) and (**B**) square waveform (SQ). Error bars represent standard error. (**C**) Correlation between exponential and square waveform.

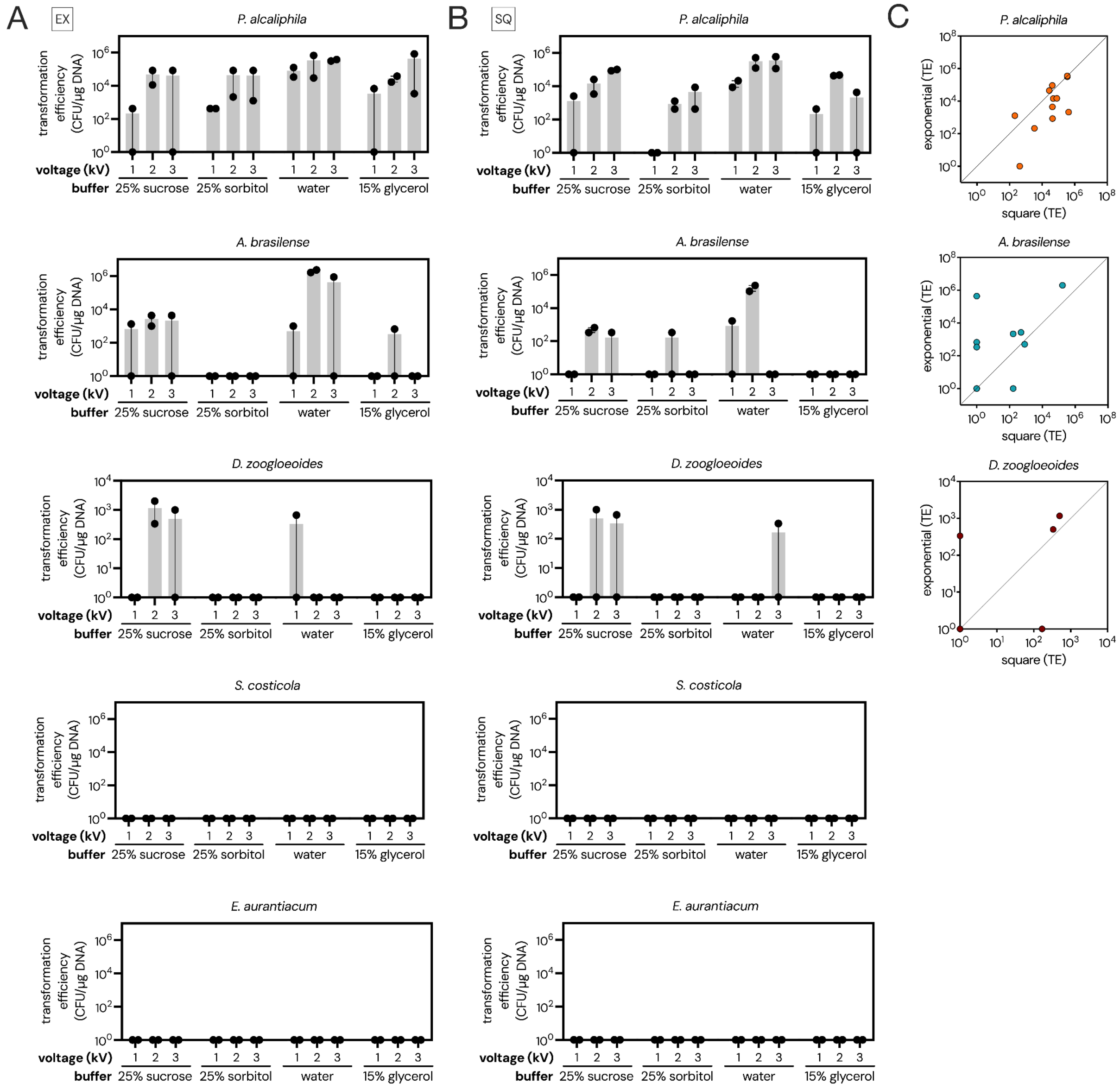

**Supplementary Figure 15. Solid selection for 24-condition electroporation screen using POSSUM plasmid library delivery to five bacteria.** Transformation efficiency as measured on solid media following plasmid library delivery to *P. alcaliphila*, *A. brasilense*, *D. zoogloeoides*, *S. costicola*, and *E. aurantiacum*. Data are the average of two biological replicates. (**A-B**) Transformation efficiency using (**A**) exponential decay waveform (EX) and (**B**) square waveform (SQ). Error bars represent standard error. (**C**) Correlation between exponential and square waveform. TE, transformation efficiency reported as CFU/µg DNA.

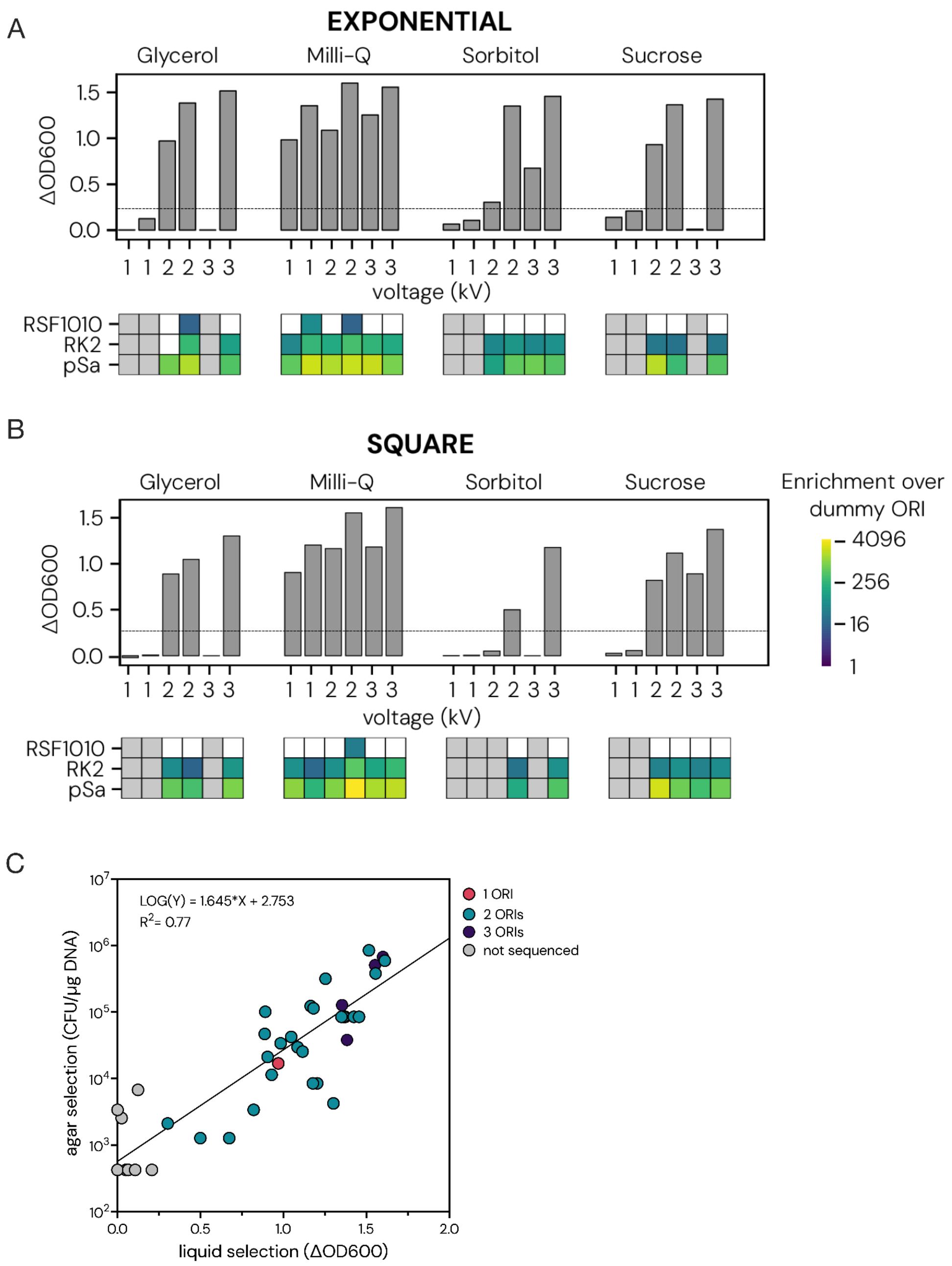


**Supplementary Figure 16. *P. alcaliphila* plasmid library ORI sequencing landscape.** (**A-B**) Results of liquid selection following POSSUM plasmid library delivery (shown as $\Delta OD_{600}$) using (**A**) exponential decay and (**B**) square waveform. For each voltage tested, data representing two biological replicates are displayed side by side on the X-axis. The dashed line indicates the sequencing cutoff, where any sample >0.25 $\Delta OD_{600}$ was processed for ORI analysis. ORI heatmap (below graphs) displays which ORI was identified in each sample. Color gradient indicates fold-enrichment of each ORI over dummy control in sequencing samples. Gray boxes indicate the sample was not sequenced, white boxes indicate that the ORI was not detected. (**C**) Correlation between liquid ($\Delta OD_{600}$) and solid (transformation efficiency reported as CFU/µg DNA) selection for all *P. alcaliphila* screen results. Color indicates the number of ORIs detected in the sample.

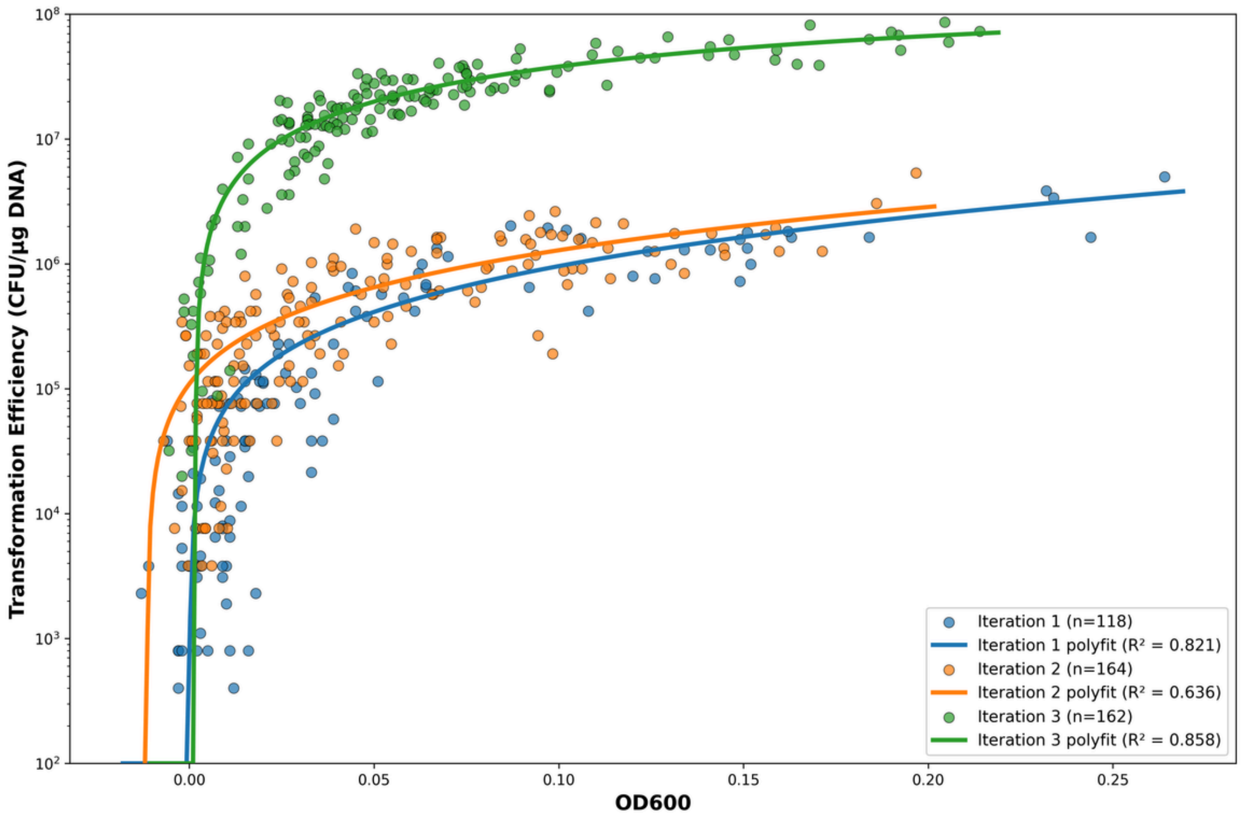


**Supplementary Figure 17. Correlation between solid and liquid selection for active learning.** $OD_{600}$ of transformants selected in liquid media compared to transformation efficiency obtained from solid selection for three cycles of active learning.

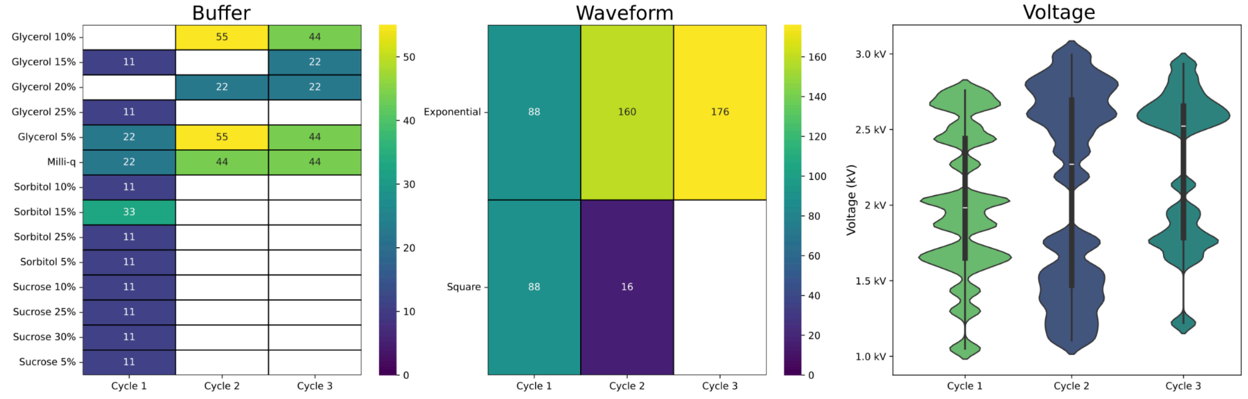


**Supplementary Figure 18. Buffer, waveform, and voltage conditions tested during three cycles of active learning.** The number of wells tested for each buffer condition (left) and each waveform (center) in each cycle of active learning for *C. necator* is shown. Empty values had 0 samples tested. Right: the distribution of voltages tested in each cycle of active learning.

### Supplementary Tables

#### Supplementary Table 1. Plasmids and plasmid libraries used in this study.

| Plasmid or library name | Origin of replication | Selective marker | Addgene ID | Source |
|---|---|---|---|---|
| Individual plasmids | | | | |
| pAKgfp1 | pBBR1 | ampicillin | 14076 | [1] |
| pAKgfp1-kan | pBBR1 | kanamycin | n/a | This study |
| pGLDV_5 [28o1G] | ColE1 | kanamycin | 199065 | [2] |
| pGL2_168 | RSF1010 | gentamicin | 199100 | [2] |
| pGL2_170 | pSa | gentamicin | 199102 | [2] |
| pGL2_270 | pSa | kanamycin | n/a | This study |
| 4-plasmid pool (gentamicin) | | | | |
| pGL2_166 | RK2 | gentamicin | 199098 | [2] |
| pGL2_168 | RSF1010 | gentamicin | 199100 | [2] |
| pGL2_174 | pNG2 | gentamicin | 199106 | [2] |
| pGL2_176 | p15A | gentamicin | 199108 | [2] |
| 17-plasmid pool (gentamicin) | | | | |
| pGL2_161 | pAL5000 | gentamicin | 199093 | [2] |
| pGL2_162 | pUB110 | gentamicin | 199094 | [2] |
| pGL2_163 | pAMβ1 | gentamicin | 199095 | [2] |
| pGL2_164 | pIP501 | gentamicin | 199096 | [2] |
| pGL2_165 | pSC101ts | gentamicin | 199097 | [2] |
| pGL2_166 | RK2 | gentamicin | 199098 | [2] |
| pGL2_167 | pBBR1 | gentamicin | 199099 | [2] |
| pGL2_168 | RSF1010 | gentamicin | 199100 | [2] |
| pGL2_169 | pIJ101 | gentamicin | 199101 | [2] |
| pGL2_170 | pSa | gentamicin | 199102 | [2] |
| pGL2_171 | pWKS1 | gentamicin | 199103 | [2] |
| pGL2_172 | pSG5 | gentamicin | 199104 | [2] |
| pGL2_173 | pBBR1-UP | gentamicin | 199105 | [2] |
| pGL2_174 | pNG2 | gentamicin | 199106 | [2] |
| pGL2_175 | Dummy/Control | gentamicin | 199107 | [2] |
| pGL2_176 | p15A | gentamicin | 199108 | [2] |
| pGL2_177 | PBC1 | gentamicin | 199109 | [2] |
| 10-plasmid pool (chloramphenicol) | | | | |
| pGL2_231 | Dummy/Control | chloramphenicol | n/a | This study |
| pGL2_238 | pUB110 | chloramphenicol | n/a | This study |
| pGL2_239 | pAMβ1 | chloramphenicol | n/a | This study |
| pGL2_240 | pBC1 | chloramphenicol | n/a | This study |
| pGL2_241 | pBP1 | chloramphenicol | n/a | This study |
| pGL2_242 | pCB102 | chloramphenicol | n/a | This study |
| pGL2_243 | pCD6 | chloramphenicol | n/a | This study |
| pGL2_244 | pIM13 | chloramphenicol | n/a | This study |
| pGL2_245 | pMB1 | chloramphenicol | n/a | This study |
| pGL2_246 | pTHT15 | chloramphenicol | n/a | This study |

**Supplementary Table 2. Strains used in this study.**

| Strain | Phylum | Family | Gram | CVM ID | Source & Catalog # |
|---|---|---|---|---|---|
| *Azospirillum brasilense* Sp 7 | Pseudomonadota | Azospirillaceae | Negative | CVM081 | ATCC 29145 |
| *Cupriavidus necator* 337 / H16 | Pseudomonadota | Burkholderiaceae | Negative | CVM130 | ATCC 17699 |
| *Duganella zoogloeoides* OSU 115 | Pseudomonadota | Oxalobacteraceae | Negative | CVM038 | ATCC 25935 |
| *Escherichia coli* MG1655 | Pseudomonadota | Enterobacteriaceae | Negative | CVM005 | ATCC 700926 |
| *Exiguobacterium aurantiacum* Colo. Road | Bacillota | Bacillaceae | Positive | CVM028 | ATCC BAA-333 |
| *Halomonas elongata* 1 H 9 | Pseudomonadota | Halomonadaceae | Negative | CVM024 | ATCC 33173 |
| *Piscinibacter sakaiensis* 201-F6 | Pseudomonadota | Sphaerotilaceae | Negative | CVM085 | DSMZ 112585 |
| *Pseudomonas alcaliphila* IAM 14884 | Pseudomonadota | Pseudomonadaceae | Negative | CVM010 | ATCC BAA-571 |
| *Salinivibrio costicola* 18AG | Pseudomonadota | Vibrionaceae | Negative | CVM029 | ATCC BAA-952 |
| *Shewanella amazonensis* SB2B | Pseudomonadota | Shewanellaceae | Negative | CVM030 | ATCC 700329 |
| *Shewanella indica* ACDC | Pseudomonadota | Shewanellaceae | Negative | CVM016 | ATCC BAA-2732 |
| *Shewanella oneidensis* MR-1 | Pseudomonadota | Shewanellaceae | Negative | CVM007 | ATCC 700550 |
| *Shewanella putrefaciens* 95 | Pseudomonadota | Shewanellaceae | Negative | CVM037 | ATCC 8071 |
| *Shewanella putrefaciens* 200 | Pseudomonadota | Shewanellaceae | Negative | CVM018 | ATCC 51753 |

**Supplementary Table 3. Previously reported electroporation conditions for strains used in this study.** EX, exponential decay; TE, transformation efficiency.

| Bacteria | Strain | Electroporation conditions | | | | | | Source |
|---|---|---|---|---|---|---|---|---|
| | | Buffer | Waveform | Voltage (kV) | Electric field (kV/cm) | Resistance (Ω) | Capacitance (μF) | |
| *E. coli* | DH5α | 10% glycerol | EX | 2.5 | 12.5 | 200 | 25 | [3] |
| | MG1655 | 10% glycerol | EX | 2.5 | 12.5 | 200 | 25 | [4] |
| *P. alcaliphila* | AL15-21$^{T}$ | 0.5 M sucrose | EX | 1.25 | 12.5 | 200 | 25 | [5] |
| *S. amazonensis* | SB2B | 1 M sorbitol | EX | 1.2 | 12 | 600 | 10 | [6] |
| *C. necator* | H16 | 50 mM $CaCl_2$ wash<br>0.2 M sucrose | EX | 1.15 | 11.5 | 200 | 25 | [7] |
| *S. oneidensis* | MR-1 | 10% glycerol | EX | 1.2 | 12 | 600 | 10 | [8] |
| | | 1 M sorbitol | EX | 1.2 | 12 | 600 | 10 | [6] |
| *A. brasilense* | Sp 7 | 10% glycerol | EX | 1.5 | 7.5 | 400 | 25 | [9] |

**Supplementary Table 4. Primers used in this study.** Regions of homology to the fragment being amplified are shown in bold. The underlined regions in PCR 1 primers represent 5' primer extensions which introduce stub sequences enabling binding of dual indexed Illumina i5 and i7 primers in the subsequent PCR 2 step.

| Primer | Sequence | Description |
|---|---|---|
| pAKgfp1_FWD | TGCTCGATGAGTTTTTCTAA**ACTGTCAGACCAAGTTTACT** | Forward primer for pAKgfp1 backbone amplification for Gibson assembly of pAKgfp1-kan |
| pAKgfp1_REV | TCAGAGATTTTGAGACACAA**GGCAAATATTATACGCAAGGC** | Reverse primer for pAKgfp1 backbone amplification for Gibson assembly of pAKgfp1-kan |
| pGLDV_5[28o1G] FWD | CCTTGCGTATAATATTTGCC**TTGTGTCTCAAAATCTCTGATG** | Forward primer for pGLDV_5[28o1G] KAN marker amplification for Gibson assembly of pAKgfp1-kan |
| pGLDV_5[28o1G] REV | AGTAAACTTGGTCTGACAGT**TTAGAAAAACTCATCGAGCATC** | Reverse primer for pGLDV_5[28o1G] KAN marker amplification for Gibson assembly of pAKgfp1-kan |
| oCG0038 | TCCCTACACGACGCTCTTCCGATCT**GGGTCACGCGTAGGACG** | PCR 1 forward primer for amplification of the plasmid barcode |
| oCG0039 | GAGTTCAGACGTGTGCTCTTCCGATCT**CCAGCTTCACACGGCGT** | PCR 1 reverse primer for amplification of the plasmid barcode |
| T_UDI_i501 | AATGATACGGCGACCACCGAGATCTACACAAGCTATAGCACACTCTT**TCCCTACACGACGCTCTTCCGATCT** | PCR 2 forward example, used to add Illumina i5 index to the sample |
| T_UDI_i701 | CAAGCAGAAGACGGCATACGAGATTTAACCGCGCGTGACTG**GAGTTCAGACGTGTGCTCTTCCGATCT** | PCR 2 reverse example, used to add Illumina i7 index to the sample |

**Supplementary Table 5. Electroporation parameter constraints for Bayesian optimization algorithm.**

| Parameter Type | Varies By | Parameter | Range | Increments |
|---|---|---|---|---|
| Electrical | Column (1–12) | Voltage | 1000 - 3000 V | 5 V |
| | | Waveform | Square wave (SQ), exponential decay (EX) | n/a |
| | | Capacitance (EX) | 10, 25, 35, 50, 60 µF | n/a |
| | | Resistance (EX) | 150 - 600 Ω | 25 Ω |
| | | Pulse Duration (SQ) | 10 µs - 1000 µs | 5 µs |
| Chemical | Row (A–H) | Buffer | Sucrose, Sorbitol, Water, Glycerol | n/a |
| | | Buffer Concentration | 5 - 30%, *Water is always 0% | 5% |

**Supplementary Table 6. Growth conditions and selective antibiotic concentrations used for bacteria in this study.**

| Strain | Media | Temperature (°C) | Antibiotic | Concentration (μg/mL) |
|---|---|---|---|---|
| *A. brasilense* | TSB | 30 | Gentamicin | 20, 40 |
| *D. zoogloeoides* | YENB | 30 | Gentamicin | 20 |
| *C. necator* | NB or SOB | 30 | Kanamycin | 50 |
| *E. aurantiacum* | YEMED pH 10 | 37 | Chloramphenicol | 8.5 |
| *E. coli* | LB | 37 | Ampicillin<br>Kanamycin | 100<br>50 |
| *H. elongata* | LB | 30 | Gentamicin | 20 |
| *P. alcaliphila* | LB | 37 or 30 | Kanamycin | 50 |
| *P. sakaiensis* | NBRC 802 | 30 | Gentamicin | 20 |
| *S. amazonensis* | LB | 37 or 30 | Kanamycin | 50 |
| *S. costicola* | MB | 30 | Gentamicin | 80 |
| *S. indica* | LB | 30 | Gentamicin | 200 |
| *S. oneidensis* | LB | 30 | Kanamycin | 50 |
| *S. putrefaciens* 95 | LB | 30 | Gentamicin | 20 |
| *S. putrefaciens* 200 | LB | 30 | Gentamicin | 20 |

**Supplementary Table 7. POSSUM toolkit parts for construction of chloramphenicol 10-plasmid pool.** All plasmids have a chloramphenicol (CAM) resistance cassette, bacterial mScarlet, *oriT* for RK2 conjugation machinery, and a variable origin of replication (ORI). Parts have been made available through Addgene.

| Plasmid constructed for this study | Destination vector | ORI | POSSUM Parts | | | |
|---|---|---|---|---|---|---|
| | | | Barcoded ORI | CAM resistance cassette | mScarlet cassette | *oriT* |
| pGL2_231 | pGL2_212 | Dummy/Control | pGL1_46 | pGL1_89 | pGL1_50 | pGL1_49 |
| pGL2_238 | pGL2_212 | pUB110 | pGL1_2 | pGL1_89 | pGL1_50 | pGL1_49 |
| pGL2_239 | pGL2_212 | pAMβ1 | pGL1_3 | pGL1_89 | pGL1_50 | pGL1_49 |
| pGL2_240 | pGL2_212 | pBC1 | pGL1_48 | pGL1_89 | pGL1_50 | pGL1_49 |
| pGL2_241 | pGL2_212 | pBP1 | pGL1_105 | pGL1_89 | pGL1_50 | pGL1_49 |
| pGL2_242 | pGL2_212 | pCB102 | pGL1_106 | pGL1_89 | pGL1_50 | pGL1_49 |
| pGL2_243 | pGL2_212 | pCD6 | pGL1_107 | pGL1_89 | pGL1_50 | pGL1_49 |
| pGL2_244 | pGL2_212 | pIM13 | pGL1_108 | pGL1_89 | pGL1_50 | pGL1_49 |
| pGL2_245 | pGL2_212 | pMB1 | pGL1_109 | pGL1_89 | pGL1_50 | pGL1_49 |
| pGL2_246 | pGL2_212 | pTHT15 | pGL1_110 | pGL1_89 | pGL1_50 | pGL1_49 |

**Supplementary Table 8. Experimental overview of 24-condition screen.**

| Experimental step | Parameter | Value |
|---|---|---|
| Cell preparation | Growth media | LB (if applicable) or optimal for strain |
| | Reseed dilution | 1:50 (<2 h doubling time) or 1:10 (>2 h doubling time) |
| | Growth phase | Uncontrolled |
| | Cell suspension buffer | 25% sucrose, 25% sorbitol, 15% glycerol, and water |
| | Number of washes | 3 |
| | Cell concentration | Unconcentrated (1X) |
| Electroporation | Cell volume | 95 µL |
| | DNA amount | 500 ng in 5 µL |
| | Waveform | Exponential decay and Square |
| | Voltage | 1, 2, 3 kV |
| | Resistance | 200 Ω |
| | Capacitance | 25 µF |
| | Pulse duration | 5 ms (EX), 0.6 ms (SQ) |
| | Gap size | 0.2 cm |
| Recovery | Media | LB (if possible) or optimal media |
| | Volume | 1 mL |
| | Time | 20 hours |

7. Vajente M, Clerici R, Ballerstedt H, Blank LM, Schmidt S. Using Cupriavidus necator H16 to provide a roadmap for increasing electroporation efficiency in non-model bacteria. Synthetic

Biology. bioRxiv; 2024. Available:
https://www.biorxiv.org/content/10.1101/2024.05.27.596136v1.full.pdf

8. Corts AD, Thomason LC, Gill RT, Gralnick JA. A new recombineering system for precise genome-editing in Shewanella oneidensis strain MR-1 using single-stranded oligonucleotides. Sci Rep. 2019;9: 39.

9. Vande Broek A, van Gool A, Vanderleyden J. Electroporation of Azospirillum brasilense with plasmid DNA. FEMS Microbiol Lett. 1989;61: 177–181.
